# Pathway-based approach reveals differential sensitivity of glioblastoma to E2F1 inhibition

**DOI:** 10.1101/2021.06.19.449118

**Authors:** Alvaro G. Alvarado, Kaleab Tessema, Sree Deepthi Muthukrishnan, Mackenzie Sober, Riki Kawaguchi, Aparna Bhaduri, Vivek Swarup, David A Nathanson, Steven A. Goldman, Harley I. Kornblum

## Abstract

Targeting glioblastoma (GBM) based on molecular subtyping have not yet translated into successful therapies. Here, we used gene set enrichment analysis (GSEA) to conduct an unsupervised clustering analysis to condense the gene expression data from bulk patient samples and patient-derived gliomasphere lines into new gene lists. We then identified key molecular pathways differentially regulated between tumors. These gene lists associated not only with cell cycle and stemness signatures, but also with cell-type specific markers and different cellular states of GBM. We identified the transcription factor E2F1 as a key regulator of tumor cell proliferation and self-renewal in only the subset of proliferating gliomasphere cultures predicted to be E2F1-activated and validated its functional significance in tumor formation capacity. E2F1 inhibition also sensitized E2F1-activated gliomasphere cultures to radiation treatment. Our findings indicate that a pathway-based approach can be leveraged to deconstruct inter-tumoral heterogeneity and uncover key therapeutic vulnerabilities for targeting GBM.

## Introduction

Glioblastoma (GBM) is incurable, with an overall median survival of about 14 months[1] despite maximal surgical resection, radiation, and chemotherapy with temozolomide[2]. The past decade has seen a revolution in the understanding of GBM, and studies of patient samples based on gene expression and oncogenic mutations have revealed that GBM can be parsed into distinct molecular categories--namely classical, mesenchymal, and proneural--and subsequently IDH mutated tumors[3-8]. While these classification schemes have shown some relationship to prognosis, they have largely failed to provide new therapeutic approaches.

The driver mutations of GBM result in the activation of many well-known oncogenic pathways, including the PI3K/AKT, MAPK/ERK, Rb, and mTOR pathways[9]. However, the use of pathway-specific inhibitors has not yet resulted in effective therapies. One potential explanation for this lack of efficacy is that tumors are comprised of multiple cell types with different pathway dependencies. Another is that inhibition of one pathway results in the activation of another[10]. Finally, it is possible that the identification of critical pathways driving GBM progression and recurrence is not yet complete. The prioritization of key pathways falls short of what would be required for tumor eradication because the combinatorial outcome of existing mutations, and resultant dominant pathways, cannot be conclusively inferred.

The advent of relatively easy and cost-effective sequencing methods and advances in bioinformatics open the door for rapid evaluation of gene expression, identification of key molecular pathways, and characterization of different cell types within individual tumors. For instance, we now appreciate that molecular subtypes previously used in the classification of GBM[5] can be linked to cell type-specific markers with the description of cellular states[11]. Similarly, a subpopulation of tumor cells can express markers of outer radial glia and turn on developmental programs to promote invasion[12]. Yet, a comprehensive and unbiased approach to group samples based on their pathway utilization has not yet been exploited to uncover therapeutic targets. We hypothesized that, rather than examining gene expression as a whole, analyzing targetable pathways could allow for the development of patient- or tumor class-specific therapeutics or combination therapies that go beyond the traditional inhibitors that have already been developed.

In this study, we developed a bioinformatics strategy that leverages Gene Set Enrichment Analysis (GSEA) to disambiguate tumor heterogeneity in GBM **(Figure 1).** We identified pathways and processes utilized by different tumors and individual cells within the tumors, without consideration of their driver mutations. We then implemented a downstream pipeline to ascertain key genes within enriched gene sets that were further evaluated for drug and/or molecular target selection. Using bulk RNA samples and existing single cell RNA-sequencing databases, we found that tumors can be clustered according to their enrichment of canonical and oncogenic pathway gene sets and that the new gene lists we derived from enriched gene sets reveal functionally significant differences between tumors and between cells within a tumor. We then utilized gliomasphere cultures, enriched or depleted in these gene lists, to demonstrate the functional significance of our approach using the pro-proliferative transcription factor E2F1 as an example.

**Figure 1.**
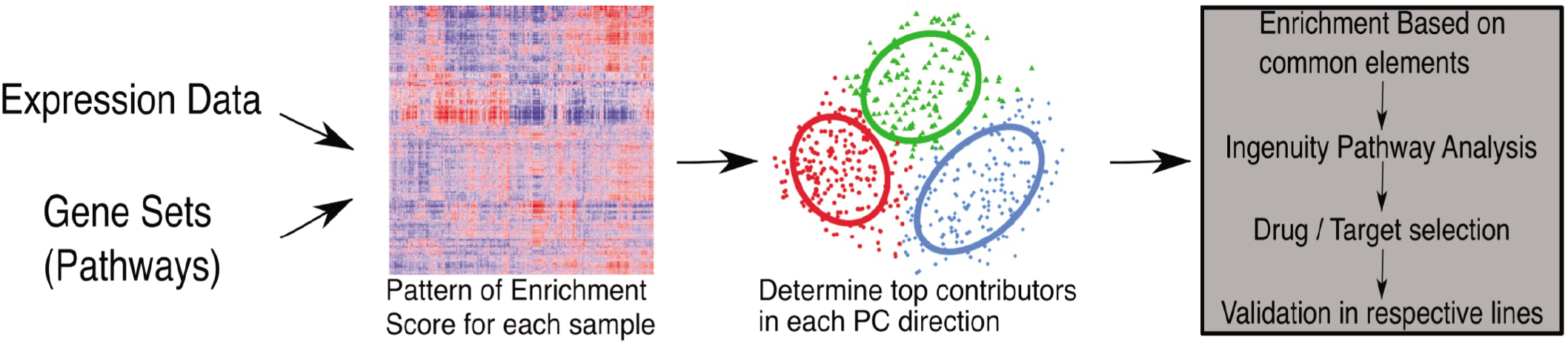
Bioinformatics pipeline for pathway-based analysis. Expression data is analyzed through gene sets to generate enrichment score profiles that are then used to cluster samples. Top contributors to particular directions are obtained and run through Ingenuity Pathway Analysis to determine targets, which can be validated in corresponding samples based on their clustering.

## Results

### GBM clustering based on molecular pathway heterogeneity is clinically significant and only modestly overlaps with prior molecular classification

To characterize intertumoral heterogeneity, we first studied presumptive pathway utilization by analyzing GBM patient samples in the Cancer Genome Atlas (TCGA[5]) using gene sets in the canonical (C2CP) and oncogenic (C6) pathway collections from the GSEA software[13, 14]. The mRNA expression of each TCGA sample was compared to the average expression of all the samples (n = 538), and normalized enrichment scores were obtained for all the gene sets comprising the two pathway collections. A heatmap representing the normalized enrichment score profile for each sample (column) demonstrates the presence of heterogeneity when either canonical **(Figure 2A)** or oncogenic **(Figure 2B)** gene sets were analyzed. We determined that 3 clusters correctly represented the data when using oncogenic and canonical pathways via non-matrix factorization and consensus clustering; the robustness of the clusters was also tested and validated using the Random Forest approach **(Supplementary Figure 1)**. We then applied principal component analysis (PCA) for dimensionality reduction to better visualize the clustering of the samples using canonical and oncogenic pathways **(Figures 2C and 2D, respectively)**. Notably, when the clustered samples were colored by their known TCGA molecular subtype, we found there was one cluster that contained most of the mesenchymal samples, while the classical and proneural subtypes were present in all clusters **(Supplementary Figure 2)**. This indicates a lack of full correlation between molecular characterization and molecular pathway expression. These findings suggest that different tumors quantitatively utilize different molecular pathways, prompting further analysis.

**Figure 2.**
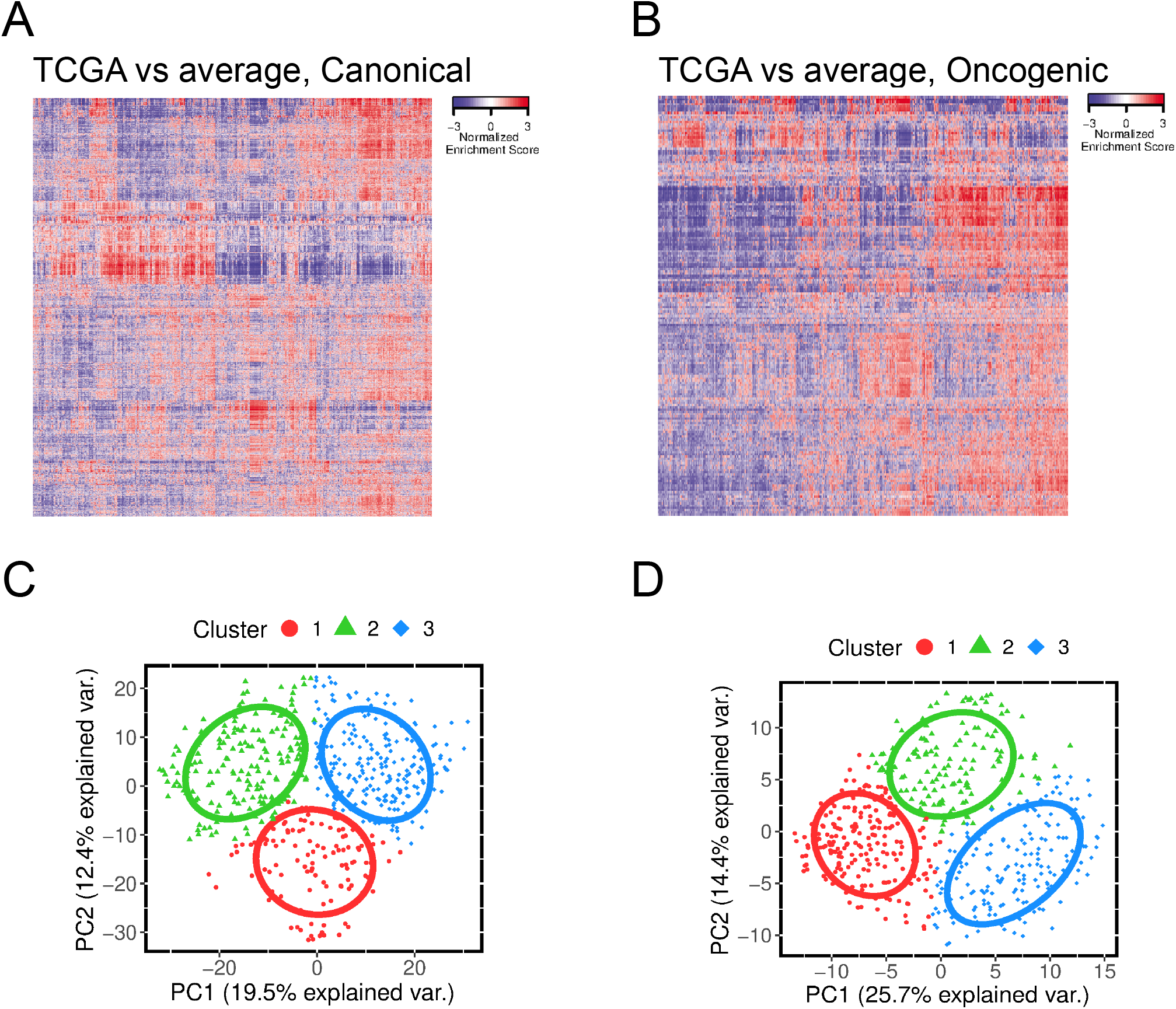
Pathway-based analysis generates three distinct clusters based on enrichment profiles. Samples from the Cancer Genome Atlas were analyzed using either canonical pathways (A) or oncogenic pathways (B) from the Gene Set Enrichment Analysis to generate heatmaps based on the enrichment profile of each sample (column) with respect to each gene set (row) in both collections. (C and D) Profiles from (A) and (B), respectively, were used to generate principal component analysis plots labeled by color and shape for each cluster. Circle lines represent the normal distribution of the samples in each cluster.

To determine whether our pathway-based classification provided functionally significant information, we examined patient survival using the pathway-based clustering information. Prior studies using TCGA groupings have found only limited association with survival, with proneural tumors having longer survival--an observation largely driven by the subset of IDH mutant tumors. As shown in **Figure 3A**, our own analysis of the TCGA categories found significant differences only between the proneural and other two groups, as previously reported [5]. However, when we utilized our new pathway-based clustering approach, we found statistically significant differences in median survival between the patients from each pair of clusters **(Figure 3B and 3C)**. For both collections of gene sets, canonical and oncogenic, the cluster with the lowest median survival was the one primarily composed of mesenchymal samples. However, approximately 30% of the samples within this cluster were characterized as classical and proneural. Similarly, using the canonical pathways collection, the cluster with the highest median survival included equal abundances of classical and proneural samples (47% each). These data challenge the idea that samples obtained from patients should be treated according to their TCGA-defined molecular phenotype and, in contrast, support the notion that tumors from different molecular backgrounds might have common signaling pathways that can be leveraged for therapeutic purposes.

**Figure 3.**
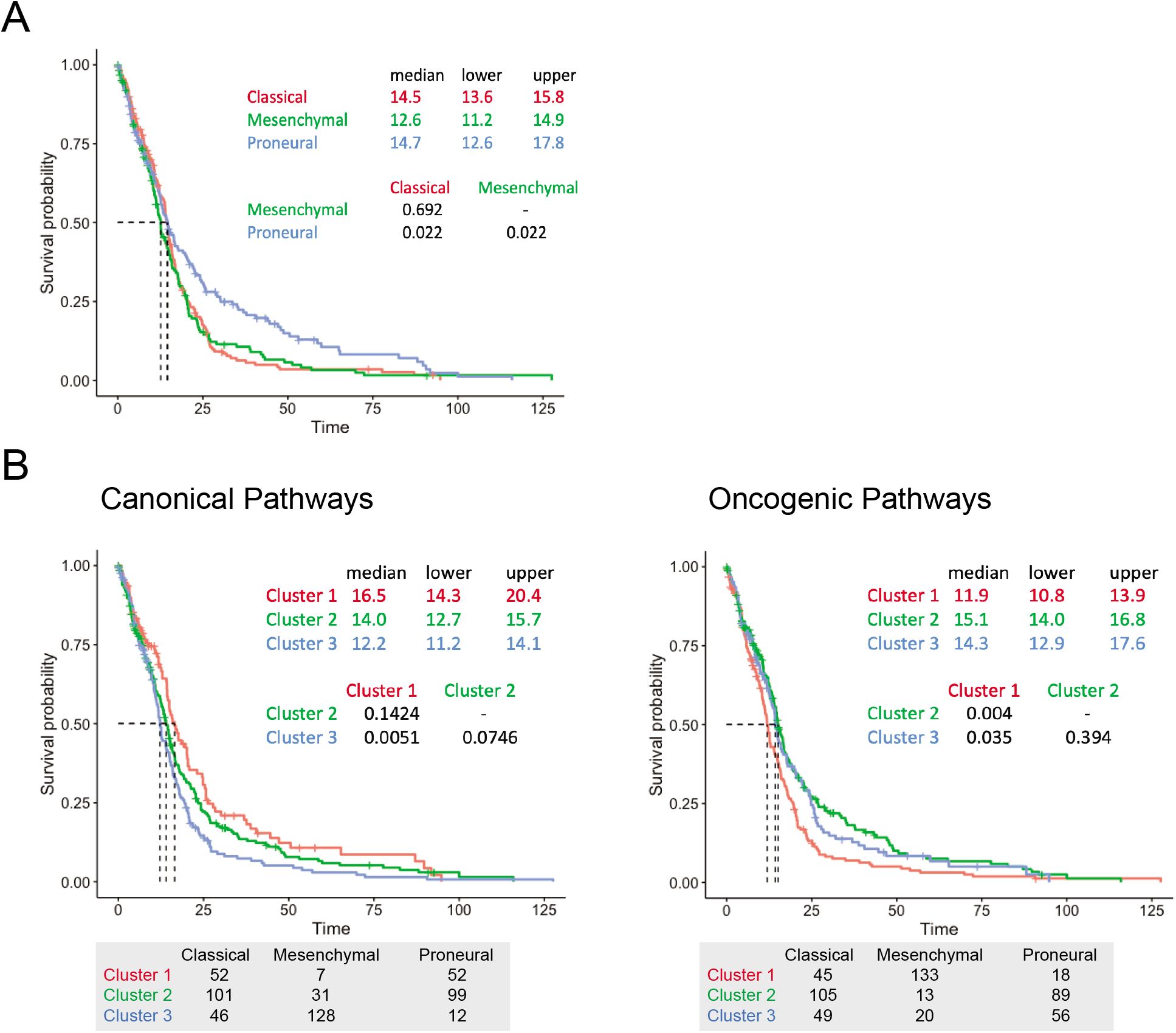
Pathway-based clusters have clinical significance and do not overlap with molecular subtypes. (A) TCGA samples were clustered based on the original molecular subtypes described, and Kaplan-Meier curves were obtained. (B and C) Samples clustered based on enrichment profiles for canonical and oncogenic gene sets, respectively, were analyzed for survival using Kaplan-Meier curves. Tables at the bottom describe the distribution of the molecular subtypes for each cluster. Dotted lines represent median survival for each curve (also described in top tables). Time shown is in months. P-values after post-hoc analyses using Bonferroni-Hochberg correction.

### GSEA gene signatures from patient-derived glioma database delineate actionable targets

Our findings thus far suggest that a pathway-based approach could be leveraged therapeutically in GBM. We recognized, however, that GSEA is an imperfect approach to assess functional pathway utilization and that individual genes or sets of genes would contribute to enrichment of multiple gene sets. Therefore, in order to more pragmatically develop potential interventions based on our analysis, we further distilled our pathway-based clustering of the whole TCGA dataset by extracting the top contributing gene sets to each principal component (PC) and direction and synthesizing their common elements into gene lists (Figure 1). We hypothesized that targeting the most highly determinative elements would allow us to target several key pathways simultaneously, even though we might be limiting our scope to shared targets and ignoring underrepresented pathways. In order to validate this approach, we performed a similar analysis on a microarray-derived database of patient-derived gliomasphere (GS) lines so that we could functionally test downstream targets. The enrichment patterns again showed heterogeneity between samples, yet they grouped into two major clusters when either canonical and oncogenic gene set collections were used **(Supplementary Figure 3A and 3B)**. This is reminiscent of a number of gene expression studies, including our own, that classify cultured glioma cells into two major groups. Gene lists were also generated for the GS dataset based on the common elements shared among the top contributing gene sets for each PC and direction, as described above for the TCGA dataset (genes in each of the gene lists generated can be found in **Supplementary Table 1**). The TCGA- and GS-based gene lists were then used to obtain enrichment scores in the gliomasphere lines **(Figure 4A)**. This dataset again separated into two main clusters in accordance with the pattern of enrichment scores generated when the oncogenic and canonical pathways were used **(Supplementary Figure 3C)**. Interestingly, there was an overlap in enrichment between some of the gene lists generated from the TCGA dataset (TCGA_C2_PC1NEG) and the gliomasphere dataset (GS_C2_PC1NEG). This was confirmed by calculating correlation values for each pair of gene lists **(Supplementary Figure 4)**. In both datasets, the strongest positive correlation was between the C2_PC1NEG lists, with coefficients of 0.78 and 0.84 in the TCGA and GS datasets, respectively. Conversely, the strongest negative correlation was between the two C2_PC1NEG lists compared with GS_C2_PC1POS (−0.57, - 0.66) in the GS dataset and with TCGA_C2_PC1POS (−0.39, −0.22) in the TCGA samples **(Supplementary Figure 4)**. These findings indicate that the identified gene lists represent independent groups of pathways for which tumor samples show differential enrichment that can potentially be exploited to uncover new therapeutic targets.

**Figure 4.**
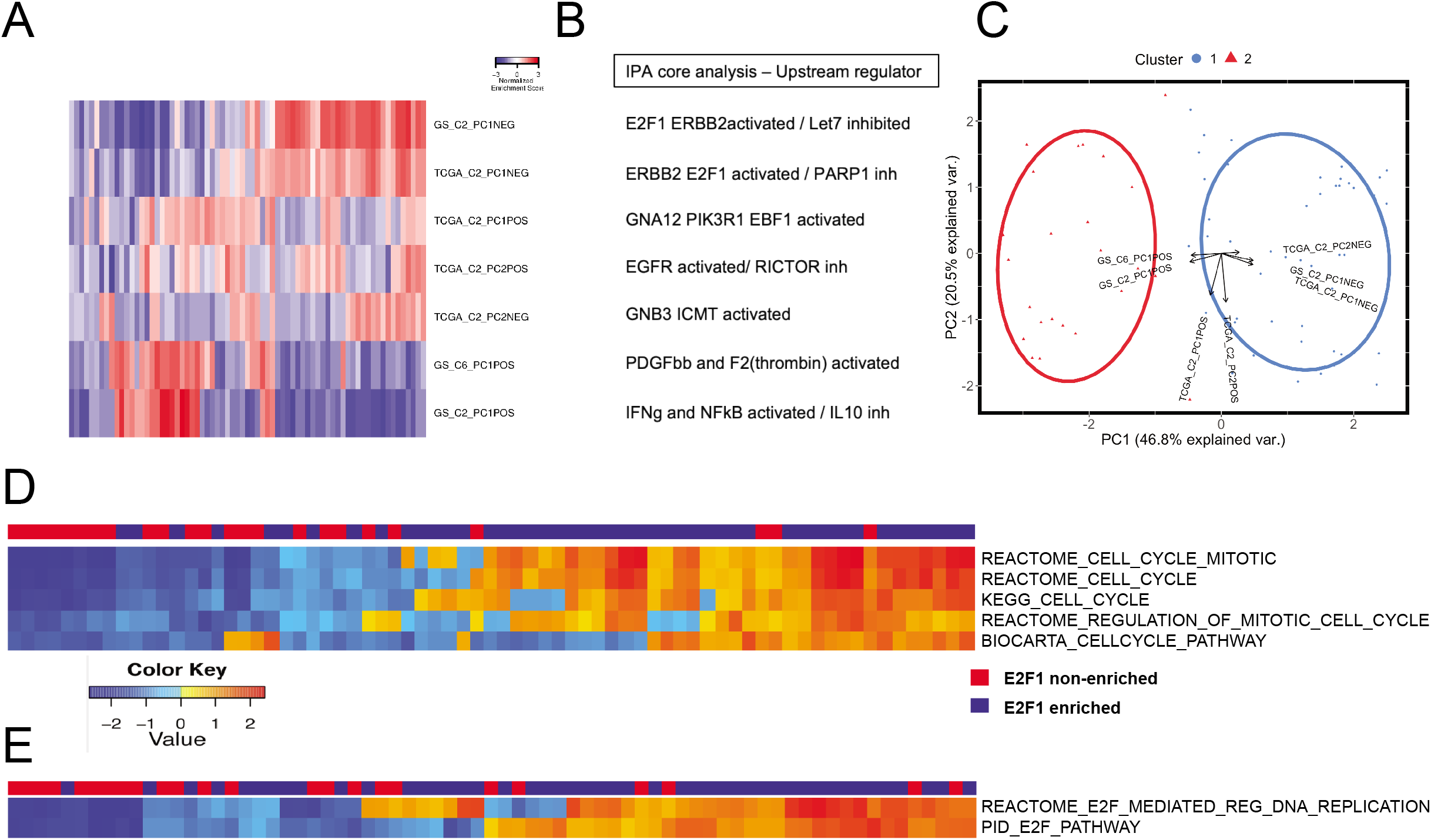
Gene lists predict E2F1 as a main target in one of the clusters found in the gliomasphere dataset. (A) Enrichment profiles using gene lists were generated for gliomasphere samples. (B) Each gene list was evaluated using IPA and top predicted activated and inhibited upstream regulators are shown. (C) PCA plot from enrichment scores generated in (A) showing how each gene list contributes to a particular direction. (D and E) Samples from both clusters were evaluated for their enrichment of cell cycle-related (D) or downstream E2F1 target (E) gene sets from the canonical pathway collection.

All the gene lists were further analyzed using Ingenuity Pathways analysis (IPA) to identify critical pathways and targets that can be leveraged therapeutically. Cluster 1 (right side of heatmap in **Figure 4A**) showed higher enrichment of the GS_C2_PC1NEG gene list and core analysis from IPA showed an increase in the expression of the E2F family of transcription factors and its downstream targets **(Figure 4B)**. Similarly, the contribution of each gene list can be appreciated in reference to both clusters **(Figure 4C)**. Activation of E2F1, together with inhibition of let7, reported to have a role in differentiation and tumor suppression [15, 16], had the most significant p-value for this particular gene list (data not shown). Since E2F1 has a known role in cell cycle progression, we examined the enrichment scores for cell cycle-related gene sets in the canonical pathways collection (C2CP). Indeed, samples that fell in the cluster with high enrichment of GS_C2_PC1NEG (and hence predicted to have activation of E2F1) had concomitant higher enrichment scores for cell cycle-related gene sets **(Figure 4D)**. Likewise, we examined the enrichment of E2F1-target gene sets from the C2CP collection and found great concordance with the samples predicted to have E2F1 activation **(Figure 4E)**. In addition, considering the correlation values between the signatures **(Supplementary Figure 4)**, we found that E2F1 activation seemed to oppose IFNγ and NFκB activation. Similarly, EGFR activation (TCGA_C2_PC2POS enrichment) appeared ubiquitously and showed correlation with most of the signatures in both datasets. These findings suggest that there are two clusters of gliomasphere samples based on their enrichment of gene lists that can be further analyzed to elucidate upstream regulators. Amongst the most meaningful differences between the samples was the fact that one cluster revealed an E2F1-activated signature exhibiting a high degree of enrichment for cell cycle and downstream target signatures.

We next re-clustered TCGA samples based on their enrichment for the gene lists described above and found 3 clusters, comparable to the original pathway analyses described in Figure 2 **(Supplementary Figure 3D)**. The E2F1-activated signatures characterized one of the clusters, whereas the EGFR signature pointed in between two of the clusters. Additionally, we analyzed raw data from the available TCGA samples (n=160) from Broad Firehose. Differential expression analysis was performed on samples using the cluster identity from the gene signatures. These data were then used to highlight the most enriched gene ontology terms for each cluster. We found each of the 3 clusters had a defined set of GO terms: cell cycle-related, extracellular membrane and inflammation, and synapse and neurotransmitter signaling **(Supplementary Figure 3E)**. These results suggest the existence of distinct clusters that can be parsed through their potential pathway utilization, highlighted by upstream regulator enrichment. Similarly, we found a greater complexity in the TCGA dataset compared with the GS dataset, as would be expected when analyzing primary and patient-derived lines, respectively.

### Pathway analysis exposes limited intratumoral heterogeneity and underscores cell cycle signature

A key consideration in the evolution of thinking about glioma heterogeneity lies in the analysis of intratumoral heterogeneity. Several single cell RNA-sequencing studies have emphasized the finding of TCGA subtype heterogeneity within tumors—that is, tumor cells from the same tumor are often classified in different TCGA groups. To assess whether potential signaling pathway utilization is similar or different within individual tumors, we interrogated a single cell RNA-seq database derived from 6 primary GBM samples[17]. Despite the reported molecular subtype diversity in each sample of this dataset, we found that, based on our GSEA approach, most cells from an individual sample clustered together based on their tumor of origin **(Supplementary Figure 5)** when either canonical or oncogenic gene sets were used. These findings suggest that, although clear intratumoral heterogeneity in gene expression exists, there may be some rationale in targeting dominant pathways as cells from each tumor have comparable enrichment profiles that contrast with cells from other tumors.

Although cells within any one tumor exhibited similarities in their enriched pathways, we recognized that our approach would not adequately identify those pathways that did indeed differ amongst cells within an individual tumor. Therefore, we analyzed each sample separately, in which each cell’s gene expression was compared to its tumor bulk control. This analysis resulted in two clusters for each tumor. Principal component loadings were obtained in order to establish which gene sets in the collection contributed the most to the apparent spatial distribution of the gene sets in the 2-dimensional PC graph. We consistently observed a cluster of cells with a high enrichment of gene sets associated with cell cycle promotion and regulation **(Supplementary Table 2)**. In contrast, the second cluster was mainly described by gene sets associated with extracellular matrix processing and growth factor signaling pathways, suggesting that both cycling and non-cycling cells could be differentially targeted. In the cell cycle-enriched clusters, we identified genes such as cyclin dependent kinases and minichromosome maintenance complex subunits that have previously been associated with glioma progression and tumor growth [18, 19]. Similarly, all 46 subunits of the proteasome complex were highly enriched in this cluster for all samples. It remains to be seen whether targeting the latter pathways will be therapeutically relevant for GBM.

In order to obtain some biological insight into the cluster not associated with cell cycle gene sets, we interrogated the gene sets associated with it using EnrichR and IPA. For each group, we selected the top common elements based on their frequency in order to generate new lists of genes that condense the information from all the gene sets. Lists were then uploaded to EnrichR (https://amp.pharm.mssm.edu/Enrichr/) and to the IPA software to obtain transcription factors, biological processes, and upstream regulators for each **(Supplementary Table 3)**. Although there are some differences between samples, there is a common theme of proliferative signals that are achieved through different mechanisms. For instance, CLOCK (circadian clock) is involved in the maintenance of pathways critical for tumor metabolism and that are upregulated during hypoxia in glioma [20]. Other pathways of interest include those involving DNA repair, epithelial-mesenchymal transition, axon guidance, and axogenesis. There are also several transcription factors that are present in most samples, such as NUCKS1 and EGR1, both of which have been previously associated with glioma (NUCKS1 in pediatric [21] and EGR1 in adult [22]) and require further mechanistic analysis. Similarly, there is a report on a feed-forward loop between EGR1 and PDGFRA that promotes proliferation and self-renewal in GSCs [23]. These findings in the scRNA-seq database indicate that there may be a potential combinatorial approach that can be tailored for each individual tumor to effectively target both cell cycle pathways and tumor-specific activated molecules.

### Gene lists differentially associate with cell cycle and stemness programs, GBM cellular states, and cell type-specific signatures

One main focus of GBM research in the last decade has been the existence of a subset of cancer cells with activated stemness programs, namely glioma stem cells, that contribute to the malignancy of the tumor [24-27] and are refractory to therapy [28, 29]. Together with the TCGA molecular subtypes, this paradigm has resulted in the development of specific therapies aimed at targeting molecules believed to be key regulators of tumor growth and invasion. The minimal clinical success of these efforts can be attributed to the limitations of the current *in vitro* and *in vivo* models used to validate these approaches. Importantly, we routinely work with patient-derived cell lines that behave differently than tumor cells in their intact tumor microenvironment in patients. It is important to establish whether critical pathways such as stem-like programs, differentiation pathways, or cell-cycle related signatures are predominantly active in these cells. To that end, we evaluated the association of our pathway-based gene signatures with cell-cycle and stemness scores in the single-cell RNA-seq dataset [17]. Consistent with our GS analyses, we found that both of our signatures predicted to have E2F1 activated (C2_PC1neg) strongly correlated with cell cycle scores **(Supplementary Figure 6A)**, while other signatures showed weak or no association. Interestingly, we found two signatures (TCGA_C2_PC1neg and TCGA_C2_PC2pos) that had strong negative correlations and one with a positive correlation (TCGA_C2_PC1pos) with stemness score **(Supplementary Figure 6B)**.

The negative correlation between activation of E2F1 and stemness score is not surprising given the fact that tumor cells are believed to be in either a proliferative or a stemness state. However, EGFR activation (TCGA_C2_PC2pos) has not been linked to a decrease in stemness and compels further investigation. The gene list with positive correlation to stemness (TCGA_C2_PC1pos) has three main targets per IPA analysis. First, phosphatidylinositol 3-kinase (PIK3R1) has been associated with GBM, and there are several inhibitors developed for molecules in this signaling pathway. GNA12 encodes for the G12 alpha subunit of G proteins and is of critical importance in regulating actin cytoskeletal remodeling in cells during migration, which is critical for tumor invasion. Finally, Early B-cell Factor 1 (EBF1) has been identified as a TET2 interaction partner in IDH-mutant cancers [30]. These analyses establish a novel approach for uncovering new molecular targets based on a pathway-based approach that can be leveraged for the development of new therapies.

We next used the same scRNAseq dataset [15] and compared our gene lists to previously reported transcriptional signatures of cell types in adult cortex [31] and developing human brain [32], as well as to recent descriptions of cellular states in glioblastoma [11]. As expected, our E2F1 activated gene lists both significantly correlated with G1/S and G2/M signatures in both datasets **(Supplementary Figure 7)**. Additionally, TCGA_C2_PC1POS (PIK3R1 and EBF1 activated) significantly associated with adult astrocytic markers and the AC-like molecular state, and TCGA_C2_PC2neg (ICMT activated) significantly associated with adult OPC markers and both OPC-like and NPC1-like molecular states. Notably, ICMT is a methyl transferase necessary for the targeting of CaaX proteins, which include the Ras family, to the cell membrane [33, 34]. Ras/ERK signaling has been associated with the proneural subtype [35] and NRAS is expressed at higher levels in proneural subtype compared with mesenchymal and classical (https://gliovis.bioinfo.cnio.es/). TCGA samples classified as proneural also showed higher enrichment for genes in the NPC-like and OPC-like molecular states [11]. These data suggest a targetable mechanism (trafficking to the cell membrane via ICMT) required for a specific protein (Ras) upregulated in a TCGA subtype (proneural) that has correlates in GBM molecular states (OPC-like, NPC1-like). Finally, we also found a significant correlation between GS_C2_PC1pos (IFNγ and NFκB activated) with both MES1-like and MES2-like states as well as adult endothelial and mural cell markers. The latter include blood vessel-associated cell types such as pericytes and vascular smooth muscle cells. This association relates to the extraordinary plasticity of glioma cells in response to their microenvironment. These data suggest IFNγ and NFκB pathways are activated in cells in the mesenchymal states that undergo vascular mimicry and express markers related to endothelial cells and pericytes that have been associated with tumor progression and recurrence[36-38]. Of note, IL-10 is also predicted to be inhibited in this gene list; given IL-10 is an anti-inflammatory cytokine, this suggests an inflammatory microenvironment would promote this particular molecular state.

### Weighted gene co-expression network analysis reveals distinct regulatory modules in each cluster

To further determine the biological relevance of the clusters identified in our GS dataset, we performed differential expression analysis and weighted gene co-expression network analysis (WGCNA). Gene ontology confirmed the enrichment of cell cycle signatures in the E2F1-enriched cluster **(Figure 5A)**. Likewise, WGCNA identified 26 modules **(Figure 5B)** that were distinctly associated with one of the two clusters **(Figure 5C)**. We took the top 3 enriched modules for each cluster and performed gene ontology enrichment analysis. Blue, brown, and light cyan modules (associated with E2F1 activation) showed enrichment of cell cycle, cell division, and DNA replication, in addition to processes associated with neurogenesis, neuron differentiation, and gliogenesis **(Supplementary Table 4)**. Moreover, two of the three modules also showed enrichment in their promoter region for E2F1 and other members of the E2F family. Conversely, black, green, and magenta modules (associated with the non-E2F1 enriched cluster) showed enrichment for inflammatory response, cell migration, and chemotaxis, as well as immune response, angiogenesis, and regulation of apoptotic processes. In these modules, we found genes with enrichment for transcription factors involved in inflammatory responses, such as CEBPB and the interferon regulatory transcription factor (IRF) family, as well as C2H2 zinc finger family members, including EGR1 and SP/KLF, which regulate proliferation, differentiation, and apoptosis cellular processes. We similarly identified the hub genes in each module, determined by how associated they are to the other members of their module[39], and colored them by module name in **Figure 5B**. Like the modules as a whole, the hub genes had different characteristics for modules associated with the E2F1 activated and non-activated clusters. Namely, the enriched E2F1-related modules show presence of known stem cell markers SOX2 and OLIG2, associated with self-renewal and persistent proliferation, as well as markers of cell division, like PLK4. Conversely, the hub genes of the other set of modules include IL-8, IL-6, and other inflammatory cytokines, together with CD44, which has been associated with a more invasive phenotype.

**Figure 5.**
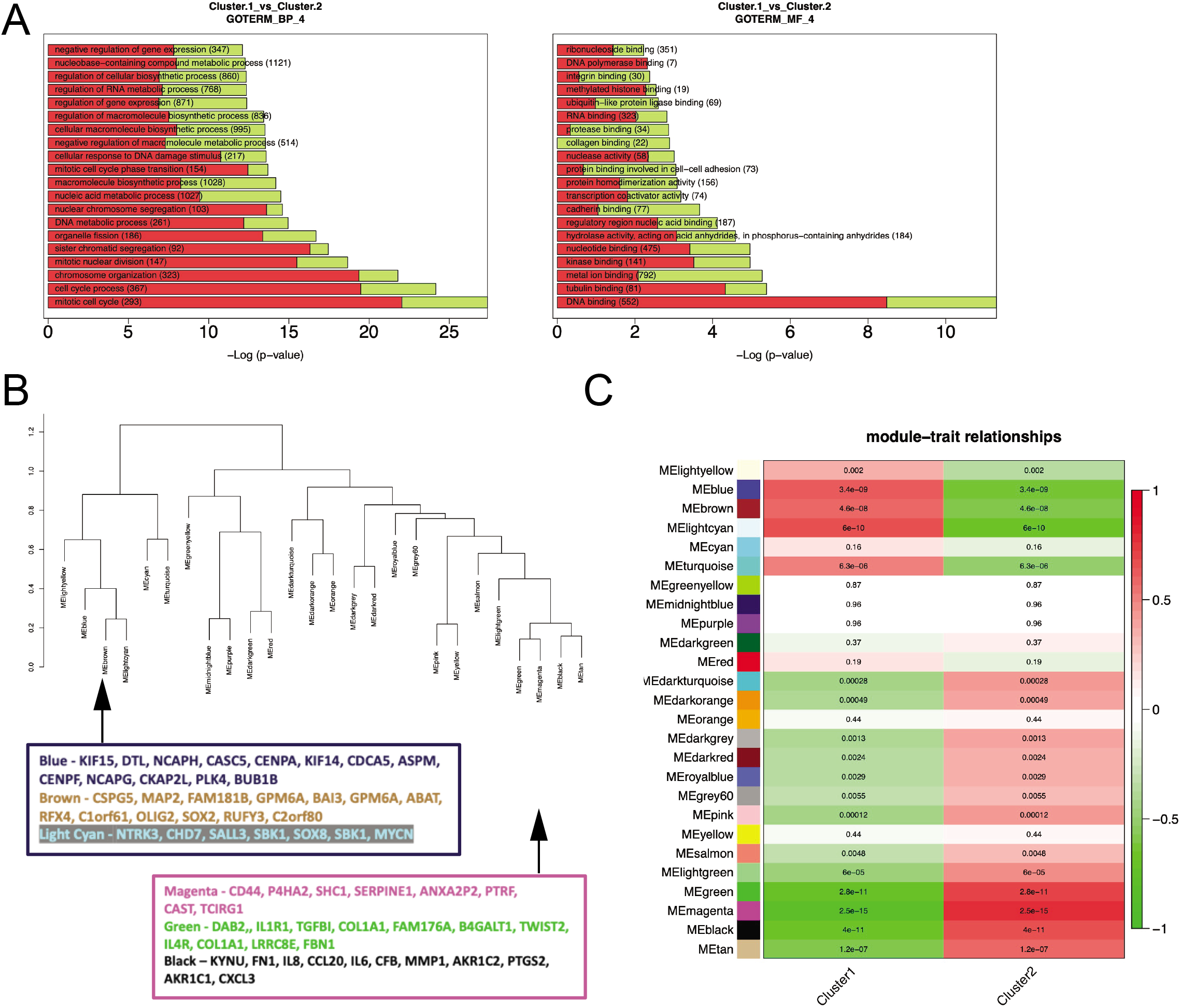
Differential expression analysis and weighted gene co-expression network analysis underscores cell cycle enrichment and reveals distinct modules in both clusters. (A) Gene expression from the gliomasphere dataset was assessed through differential expression analysis. Cluster 1 represents samples in the E2F1-enriched cluster. Two examples from Gene Ontology are shown, depicting high enrichment for cell cycle terms. (B) WGCNA generated 26 modules when samples were analyzed based on their enrichment profiles for the gene lists. (C) Modules are ranked based on their abundance in both clusters. Modules at the top are highly enriched in the E2F1-activated cluster (cluster 1).

From a clinical perspective, we wanted to know if the hub genes would be viable as therapeutic targets. To this end, we took advantage of a prior study comparing gene expression in the cellular fraction containing tumor initiating cells, termed glioma-derived progenitors cells (GPCs) and normal, non-transformed glial progenitor cells (nGPCS) [40]. For the E2F1-related modules, the blue module had several hub genes (DTL, CASC5, CDCA5, ASPM, CENPF, BUB1B) whose expression was at least 4-fold change higher in glial GPCs. Similarly, MYC in the light cyan module was 12-fold more highly expressed in glial GPCs relative to nGPCs. Evaluation of the non-E2F1 associated hub genes uncovered that CD44 and COL1A1, both associated with invasion, were highly expressed in glial GPCS (over 22-fold change higher related to nGPCs). Two other genes associated with migration and invasion, FN1 and SERPINE1, were also at least 4-fold higher in glial GPCs. These data confirm our pathway-based analyses that generated two clusters characterized by cell cycle enrichment contrasted with inflammatory and promotion of invasion pathways that are predicted to have lower chances of off-target effects based on their expression profiles on glioma-derived and normal progenitor cells.

### Differential effects of E2F1 silencing and candidate therapeutics support the functional significance of pathway-based heterogeneity

Our findings of potentially differential dependence on E2F1 and its downstream targets in highly proliferative gliomasphere cultures was somewhat surprising, as this transcription factor is often thought to be primarily involved in proliferation and cell cycle regulation. To investigate this further and to validate our general approach, we used silencing technology to evaluate the cellular effects of E2F1 suppression in samples from E2F1-activated and non-activated clusters. HK217 and HK301, members of the E2F1-activated cluster, showed a marked decrease in stem (sphere-forming) cell frequency in a limiting dilution assay (LDA) in cells with E2F1 knockdown compared with control **(Figure 6A)**. These effects were not observed in HK357 or HK408, lines that were not enriched for an E2F1-activated signature (Figure 6A). Likewise, knockdown of E2F1 resulted in compromised overall cell proliferation in E2F1-enriched samples when E2F1 expression was suppressed **(Figure 6B)**, compared with non-enriched cells where E2F1 knockdown did not significantly alter proliferation. In order to determine whether there are potential pharmaceuticals that can target the gliomasphere clusters with and without an E2F1-activated signature, we utilized the drug predictive upstream tool in IPA. We found that the gene lists that had E2F1 as a common element shared fulvestrant and calcitriol as very significant hits. Samples from the E2F1-enriched cluster, HK217 and HK336, were treated with these drugs and assessed for sphere formation capacity. HK217 showed a significant decrease with both treatments, and HK336 showed a significant response with calcitriol treatment **(Figure 6C)**. Conversely, HK408 (from the non E2F1-enriched cluster) did not show a significant change in sphere forming capacity after either treatment (data not shown).

**Figure 6.**
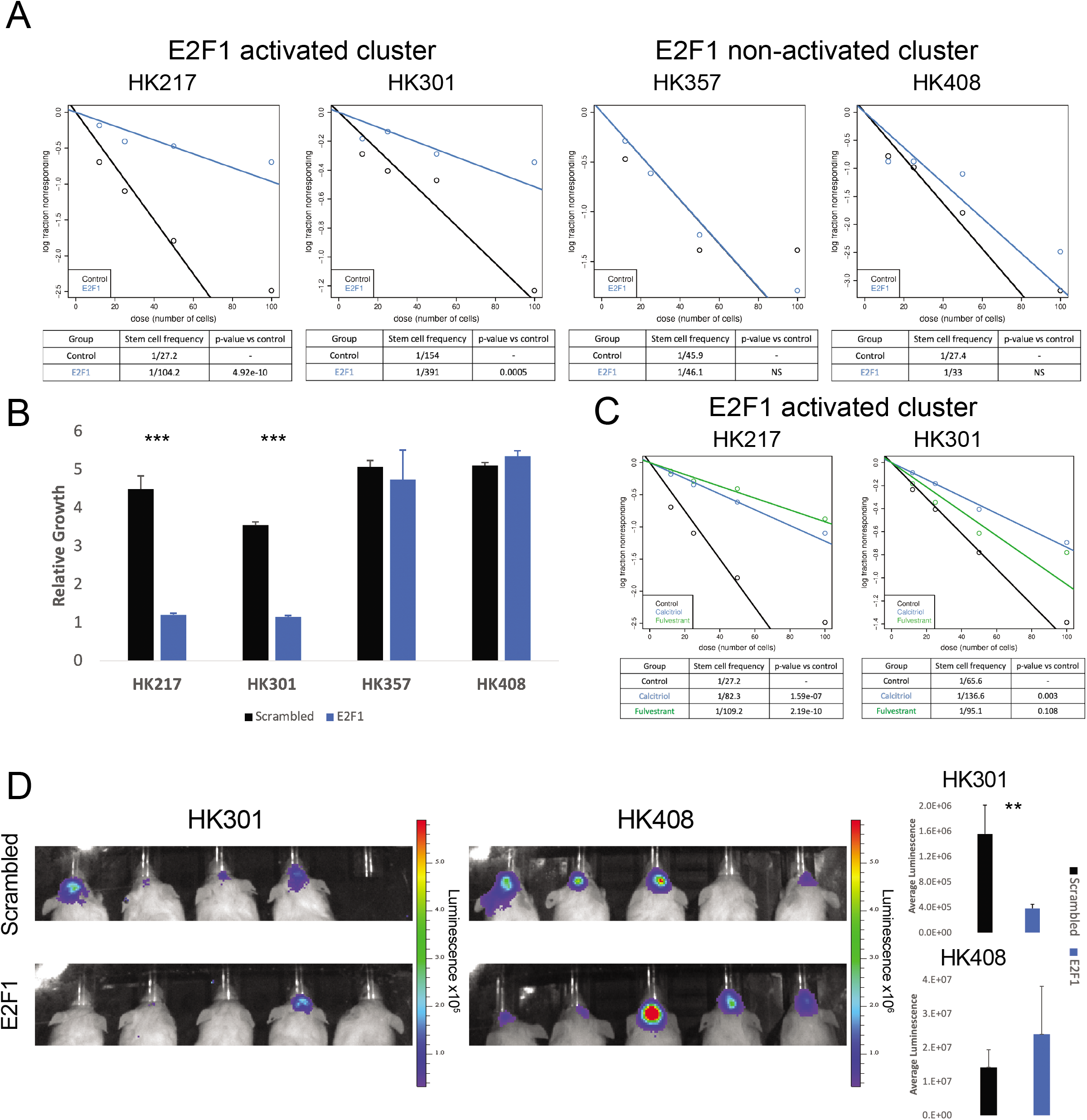
E2F1 silencing compromises self-renewal and proliferation *in vitro* and tumor formation *in vivo*. (A) Samples from both clusters were treated with control (scrambled) or E2F1 siRNA and plated under limiting dilution in a 96-well plate. Graphs depict the number of wells that did not form spheres after 10 days vs. the number of cells plated (a vertical line implies all wells formed spheres). (B) Cells treated with scrambled or E2F1 siRNA were plated at a density of 2,000 cells per well in a 96-well plate in quadruplicate, and their growth was evaluated using luminescence. Relative growth is the fold change compared to basal measurement. (C) Cells from the E2F1-activated cluster were treated with two drugs identified using IPA, and their sphere formation capacity was evaluated after 10 days. (D) Cells treated with scrambled or E2F1 shRNA were intracranially injected in NSG mice. Luminescence was assessed four weeks after transplantation. Quantification for each group is shown on the right. Experiments in (A), (B), and (C) were performed at least three times. Data are represented as mean +/- SEM. **p<0.01 and ***p<0.001 as assessed by one-way ANOVA.

To determine the *in vivo* relevance of our findings, we assessed tumor formation capacity in HK301 (E2F1-activated) and HK408 (E2F1-non-activated) cell transduced with E2F1 KD (knockdown) and control (scrambled) lentivirus. *In vivo* bioluminescent images showed tumor growth in all mice intracranially transplanted with HK408 in both control and KD groups. Conversely, HK301 E2F1 KD cells showed limited tumor formation *in vivo* while animals injected with HK301 control cells showed tumors in four out of five mice **(Figure 6D)**. These data demonstrate that the targets uncovered by this pipeline have functional implications in patient-derived gliomaspheres.

The transcription factor E2F1 is a critical component of the cell cycle signaling machinery. However, having a cluster of samples that do not rely on this molecule for proliferative purposes, as demonstrated *in vivo* and *in vitro* by loss-of-function experiments in the non-E2F1 cluster, was surprising. We therefore decided to investigate whether E2F1 might be regulating other cellular processes in GBM. E2F1 has been demonstrated to have a role in the suppression of senescence in prostate cancer cells and proposed to be a key factor for the progression of tumors in the presence or absence of p53 or retinoblastoma[41]. Similarly, it has been described in breast cancer cells that senescence sensitivity is regulated by an interaction between E2F1 and cellular inhibitor of PP2A (CIP2A)[42]. Accordingly, we found increased expression of CIP2A in the E2F1-activated compared with the non-enriched cluster in the GS dataset **(Figure 7A)**. Correlation analyses also found highly significant correlations in both the TCGA and GS datasets between E2F1 and CIP2A expression **(Figure 7B)**. Of note, E2F1 knockdown affected CIP2A expression only in E2F1-enriched samples **(Figure 7C)**, which suggests a unique regulatory network primarily utilized in cells belonging to this cluster. We decided to test this functionally by treating cells with irradiation and measuring their capacity to resolve DNA damage as measured by γH2AX staining. HK217 and HK408 control and E2F1 KD cells were irradiated with a single dose of 8Gy, and cells were stained 12 hours later. HK408 showed comparable levels of H2AX-positive cells under both conditions (control = 80%, KD = 78%, n.s.), whereas E2F1 KD significantly impacted the capacity of HK217 cells to resolve DNA damage (control = 45%, KD = 68%, p-value = 0.01) **(Figure 7D)**. Additionally, we evaluated the enrichment of DNA repair- and senescence-related gene sets in samples from both clusters. As expected, samples in the E2F1-activated cluster showed higher enrichment for both groups of gene sets **(Supplementary Figure 8)**. These data further confirm that the pathway-based approach we have implemented in these studies has identified a specific molecular target for a cluster of samples that has both biological significance and therapeutic potential to advance treatment for patients with GBM.

**Figure 7.**
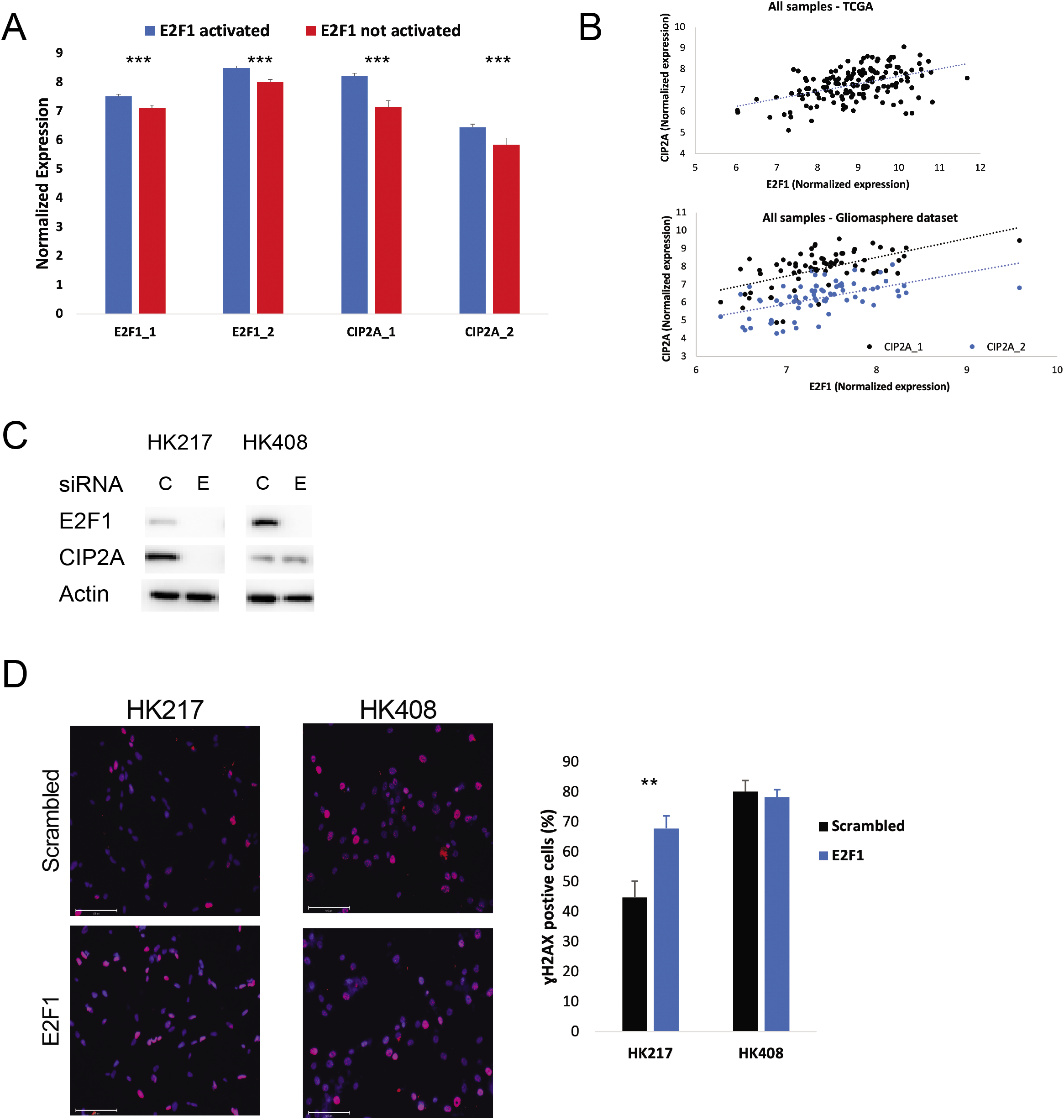
E2F1 silencing compromises DNA damage response induced after irradiation. (A) Gene expression levels of *E2F1* and *CIP2A* in the gliomasphere dataset. (B) TCGA (top) and gliomasphere (bottom) samples were plotted based on their expression levels of *E2F1* and *CIP2A* (two different probes in the gliomasphere dataset). Trend lines show significant positive correlation > 0.5. (C) Cells were treated with either C (control) or E (E2F1) shRNA, and protein was assessed for E2F1 and CIP2A levels. Actin serves as the loading control. (D) Treated cells were subjected to irradiation (8 Gy) and fixed after 12 hours for γH2AX staining (red). Nuclei were counterstained using DAPI. Quantification for each group is shown on the right. Experiments in (C) and (D) were performed at least two times. Data are represented as mean +/- SEM. **p<0.01 and ***p<0.001 as assessed by one-way ANOVA.

## Discussion

In this study, rather than focusing on driver mutations themselves, our goal was to focus on their impact on gene expression and to use the latter in an unbiased manner to assay molecular pathways that will influence the biology of the tumor. Our assumption is that while individual mutations may influence one of a number of different processes, ranging from protein phosphorylation to chromatin modifications, mutations will ultimately result in altered gene expression, which then results in modified cellular function. Although our strategy does result in a reclassification of tumors and tumor cells, the analyses described in this work present a pathway-based approach to uncover biologically relevant, actionable targets derived from the heterogenous biology inherent to glioblastoma.

GSEA is a powerful tool used to group sets of genes in a functionally relevant manner. However, the gene sets in GSEA, especially in the oncogenic collection, are often based on the responses of cells and certain tissues to genetic perturbations, and enrichment for a particular gene set does not prove that a specific process or pathway is involved. Furthermore, an individual gene or group of genes can be represented in multiple gene sets. In order to obtain more precise information from our analysis, we added an extra layer to our approach where we extracted genes that were common to multiple gene sets and that were highly associated with the directionality of the principal components. We then analyzed these “eigen-gene sets” for their enrichments in pathways and processes. This additional step allowed us to identify functionally important genes and processes.

Using our approach, gene lists were established form both bulk tumor samples and patient-derived gliomasphere datasets and associated with specific cell signatures in a single cell dataset. One of the clearest relationships we observed was the strong association between signatures associated with E2F1 activation and proliferative signatures. One potential explanation of such a finding would be that different tumors have different numbers of proliferating cells and thus differential gene expression based on their abundance. However, our findings in gliomaspheres suggest that there are more complex processes at play, as both E2F1-dependent and non-dependent cultures were highly proliferative at the time of study, indicating that the expression differences observed represented true differences in the biology of the cells. It is possible that another closely related member of the E2F family would serve the same function as E2F1 in the non-enriched population. However, one may surmise that such factors would result in similar downstream effectors and therefore would not have appeared to be enriched in our studies.

In addition to its role in proliferation, recent studies tie E2F1 function to other processes, including DNA repair. E2F1 is known to promote the expression of the DNA repair protein CIP2A, which was confirmed to be enriched in our E2F1-activated samples. Our studies confirmed inhibition of E2F1 reduced CIP2A expression and reduced the capacity to resolve irradiation-induced DNA damage in E2F1-activated gliomasphere cultures. Our analysis also identified small molecules that could selectively target this pathway and be considered for development of therapeutics in subclasses of cells. This study leveraged the availability of a large library of well-characterized gliomasphere samples. While these are thought to embody much of the complexity of GBMs, they are undoubtedly a simpler system and are likely to be enriched in actively proliferating cells, as opposed to quiescent stem or stem-like cells. Here we report differential sensitivity of tumor cells to E2F1 inhibition between distinct clusters of proliferating gliomasphere lines; yet we recognize the limitations of the gliomasphere culture system. Most studies based on expression have parsed gliomasphere cultures into two general categories, similar to our findings, rather than the multiple subtypes exhibited by tumors themselves. It is possible that other culture systems, such as organoids, would be better able to replicate the inter- and intratumoral heterogeneity observed in gliomas.

In addition to an E2F1-driven cluster of tumors and gliomaspheres, we observed clusters that were not E2F1-driven and that appeared to be more heavily reliant on other pathways. For example, the cluster on the left in Figure 4A showed diverse enrichment for gene lists whose main targets are more classical dysregulated pathways in glioblastoma, such as PIK3R1 and PDGF receptor [9]. This cluster also had a strong enrichment in the majority of its samples for an IFNγ- and NFκB-activated signature. This inflammatory and/or damage response was also observed in the TCGA dataset as one of the main components for one the clusters described in our first analysis **(Supplementary Figure 3E)**. Finally, samples predicted to have EGFR activation were equally distributed in both clusters, suggesting EGFR expression levels are not particularly informative in terms of functional diversity in glioblastoma samples.

In our single cell analysis, where we examined the pathway-based expression of all the cells within a tumor, we found that the most significantly enriched gene sets compared to the bulk tumor average were generally very similar. This suggests that the predominant pathways used by all or most of the cells within a single tumor might be targeted, but that these pathways would vary from tumor to tumor. It is important to note that individual sample analyses did separate cells from each tumor into a cell cycle-enriched cluster and non-cell cycle related cluster. In the latter, we identified EGR1 as a transcription factor upregulated in most samples **(Supplementary Table 3)**. This same molecule was underscored in the modules associated with the non-E2F1 cluster when we performed WGCNA (**Figure 5B and Supplementary Table 4**). EGR1 has been associated with O-6-Methylguanine-DNA Methyltransferase (MGMT) methylation[22]. Our findings link a potential prognostic marker to a subset of GBM samples that could have potential therapeutic implications.

Moreover, using cell signature scores rather than setting arbitrary values for cell type identities, we were able to determine some of the characteristics of individual cells within tumors. This analysis confirmed that an E2F1-driven signature correlated with genes that were related to mitosis, but inversely correlated with putative markers of stemness. It is unclear whether this is because “true” cancer stem cells are slowly dividing, or whether other factors are involved. We were able to further our correlational analysis to include cellular states described in glioblastoma [11] and cell types from normal brain development. The rationale to do these analyses was based on a recent report using scRNA-seq that uncovered a subset of glioblastoma cells with outer radial-glia signatures that were able to activate an embryonic pathway to promote invasion[12]. Our studies link cellular states to cell type-specific signatures and potential targets from our gene lists. For example, predicted ICMT activation was negatively associated with stemness and positively associated with NPC-like and OPC-like states, as well as adult OPC signatures. Similarly, IFNγ and NFκB activation were positively correlated with MES-like1 and MES-like2, as well as endothelial and mural (vasculature, pericyte) signatures. This last interaction is particularly interesting since it suggests an inflammatory environment as a driver for the expression of tumor vasculature markers. This is consistent with the capacity of glioma cells to undergo transdifferentiation into endothelial cells and pericytes to promote invasion[36-38]. These analyses provide new potential avenues for the development of innovative treatments.

Recent reports have appreciated the complexity of tumor cell expression signatures, and now the emphasis has been on providing a more holistic view of the cells within each tumor, such as cellular states [11], a single axis of variation between proneural and mesenchymal subtypes [43], and a recent report using a similar approach to ours that introduces another layer of complexity to glioblastoma heterogeneity by uncovering a mitochondrial subtype with unique vulnerabilities [44]. Our study adds to this trend by providing a novel approach to condense tumoral heterogeneity to critical gene lists that can be used to identify upstream regulators. In conclusion, we propose a combinatorial approach where precision medicine will be composed of sample-specific drugs that also provide specific vulnerabilities to be exploited with metabolic and/or immune-activating approaches. The integration of different aspects of a cell or sample is paramount for the development of new therapeutics.

## Materials and methods

### Patient and tumor datasets

Glioblastoma samples analyzed were composed of TCGA dataset [5], gliomasphere microarray dataset [45], and single cell RNAseq dataset [17] (GSE57872).

### Gene set enrichment analysis, gene list generation, and target prediction

For TCGA and gliomasphere datasets, each sample was compared to the average of the whole dataset using the canonical (C2CP) and oncogenic (C6) gene set collections from the GSEA website (http://www.gsea-msigdb.org). Enrichment profiles were then used to generate principal component analysis plots, and the contribution of each gene set to a particular direction was extracted using R-package ‘FactorMineR’. The top 20 contributing gene sets in a particular direction were compared to one another and common elements (present in at least 5) were considered for the gene lists. The names are derived from the dataset (TCGA or GS), the component (PC1 or PC2), and the direction (positive or negative). Datasets were reanalyzed with these gene lists to obtain similar clustering and for downstream analyses. Gene lists were further explored via Ingenuity Pathway Analysis using the upstream regulator tool. Targets (molecules or drugs) were predicted to be activated or inhibited for each of the gene lists used as input.

### Gene list correlation analysis

For both the TCGA and GS bulk transcriptomic datasets, Pearson correlation coefficient was computed using R (“corr.test” function in “psych” package) for each pair of gene lists using the sample-level enrichment scores previously generated. Hypothesis testing was performed for each pair to assess significance of correlation, resulting in a matrix of both raw and FDR adjusted p-values. Correlation plots were generated using the “corrplot” package in R. Displayed are the correlation coefficients for each pair and circles whose color and size reflect the coefficient value and magnitude, respectively. For pairs with non-significant correlation, the coefficients are displayed without circles. Significance was evaluated using α=0.05 for both raw (below diagonal) and FDR-adjusted (above diagonal) p-values. Correlation patterns were used to group the gene lists using hierarchical clustering, with black boxes marking the resulting clusters.

### Single cell signature score analysis

Single cell expression data (n=430) from 5 primary human glioblastoma specimens were imported from GSE57872. For each gene set of interest, single cell enrichment scores were generated as described[17]. Briefly, the enrichment score of a gene set was computed in each cell by taking the average expression of genes within the gene set and subtracting the average expression of all detected genes. Single cell enrichment scores were generated for (1) the 6 TCGA/GS gene lists discussed above, (2) the cell cycle meta-signature described in Fig 2B of [17], and (3) the stemness signature described in Table S1 of [46]. These scores were used to visualize pair-wise gene set correlations across cells, specifically between each of the 6 TCGA/GS gene lists and cell cycle (Supp Fig 5A) or stemness (Supp Fig 5B). Single cell enrichment scores were then generated for two additional groups of gene sets: developing[32] and adult[31] brain cell type markers and glioblastoma cellular state markers[11]. These scores were used in combination with the 6 TCGA/GS gene list scores to generate a correlation plot (Supp Fig 6) as described above in the “Gene list correlation analysis” section.

### Patient-derived gliomasphere cultures

Established patient-derived gliomasphere lines were cultured and maintained as previously described[47]. Experiments were performed only with lines that were cultured for less than 20 passages since their initial establishment and were tested for mycoplasma regularly.

### *In vitro* functional analysis: sphere formation and cell proliferation

Cell proliferation experiments were conducted by plating cells at a density of 2,000 cells/well in a 96-well plate in quadruplicate. Cell number was measured after 3 and 7 days and normalized to the initial reading at day 0 using the CellTiter Glo Luminescent Cell Viability Assay (Promega). The experiments shown represent fold change at day 7 relative to day 0. For sphere formation assays, cells were plated at a low density (100, 50, 25, and 12 cells per well) in 96-well plates (24 wells per density). Cells were maintained for 10 days before sphere formation was evaluated. Spheres larger than 10 cells in diameter were considered for analysis. The numbers shown represent the number of cells per well or the stem cell frequency as calculated using the Walter and Eliza Hall Institute Bioinformatics Division ELDA analyzer (http://bioinf.wehi.edu.au/software/elda/) (Hu and Smyth, 2009). All sphere formation and proliferation experiments were repeated at least three times.

### *In vivo* tumor xenografts and imaging

For tumor formation assessment, 8- to 12-week-old NOD-SCID null (NSG) mice were used in equal numbers of female and male. 5×10^4 tumor cells containing a firefly-luciferase-GFP lentiviral construct and either a scrambled or E2F1 shRNA vector were transplanted per mouse (n = 5), in accordance with UCLA-approved Institutional Animal Care and Use Committee protocols. Five mice were housed per cage, with a 12hr light/dark cycle, and were provided food and water ad libitum. Tumor growth was monitored every 2 weeks after transplantation by measuring luciferase activity using IVIS Lumina II bioluminescence imaging. ROIs were selected to include the tumor area, and radiance was used as a measure of tumor burden. Mice were monitored and sacrificed upon the development of neurological symptoms such as lethargy, ataxia, and seizures, along with weight loss and reduction in grip strength. Animals were sacrificed by CO2 asphyxiation and secondary cervical dislocation.

### Lentivirus transduction in gliomasphere lines

Lentiviral vector particles containing E2F1 and scrambled shRNAs were purchased from Abmgood. Cells were transduced with the corresponding viruses for 48 hours and selected with puromycin (Sigma). Knockdown of E2F1 was confirmed using immunoblotting in treated samples.

### Immunofluorescence analysis

Cells were plated on 24-well plates pretreated with laminin overnight. After two days of culture, cells were fixed in 4% paraformaldehyde for 15 min at room temperature, followed by blocking and overnight incubation at 4°C with γH2AX primary antibody (Cell Signaling Technology). Cells were then incubated with species-specific goat secondary antibody coupled to AlexaFluor dye (568, Invitrogen) and Hoechst dye for nuclear staining for two hours at room temperature. Plates were imaged using EVOS microscope, and quantification of positively stained cells was performed manually using ImageJ.

### Immunoblotting analysis

Cells of interest were lysed using RIPA lysis buffer (Thermo Scientific), and protein concentrations were calculated using a BCA protein assay (BioRad). After denaturation with Laemmli buffer (BioRad), 10 mg of total protein was loaded on 4-12% polyacrylamide SDS-PAGE gels (NuPage, Themo Scientific), transferred to polyvinyl difluoride (PVDF) membranes (Millipore), and probed using the following antibodies: E2F1 (Santa Cruz Biotechnology, 1:500), CIP2A (Santa Cruz Biotechnology, 1:1000), and b-Actin (Cell Signaling Technology, 1:2000) for loading control. Species-specific horseradish peroxidase (HRP)-conjugated secondary antibodies were used for detection (Cell Signaling Technology, 1:5000). Membranes were developed using ECL-2 reagent (Pierce Biotechnology). All western blots were performed at least three times.

### Irradiation of gliomaspheres

Cells were irradiated at room temperature using X-ray irradiator (Gulmay Medical Inc., Atlanta, GA) at a dose rate of 5.519 Gy/min for the time required to apply an 8Gy dose. The X-ray beam was operated at 300 kV and hardened using a 4mm Be, a 3mm Al, and a 1.5mm Cu filter, and calibrated using NIST-traceable dosimetry.

### Statistical analysis

Reported data are mean values ± standard error of the mean for experiments conducted at least three times. Unless stated otherwise, one-way ANOVA was used to calculate statistical significance, with p-values detailed in the text and figure legends. P-values less than 0.05 were considered significant. Correlation analyses were performed using Pearson coefficient. Log-rank analysis was used to determine the statistical significance of Kaplan-Meier survival curves. Data analysis was done using R v 3.6.3[48].

## Author contributions

A.G.A, S.M and M.S conducted in vitro and in vivo experiments. A.G.A, K.T, R.K, and V.S performed bioinformatics analysis. A.B and D.A.N contributed resources and reagents. A.B, V.S, D.A.N, and S.A.G provided valuable inputs in experimental design and data analysis. A.G.A and H.I.K conceptualized the study, designed the experiments and wrote the manuscript. All authors read, revised, and approved the final manuscript.

## Acknowledgements

The authors thank the members of the Kornblum lab for insightful comments. We would like to acknowledge the UCLA pathology and flow cytometry cores for their technical assistance. A.G.A was funded by the UC President’s Postdoctoral Fellowship. H.K was funded by the Dr. Miriam and Sheldon G. Adelson Medical Research Foundation, and the UCLA SPORE in Brain Cancer P50 CA211015-01A1.

## References

1. Johnson, D.R. and B.P. O’Neill, Glioblastoma survival in the United States before and during the temozolomide era. J Neurooncol, 2012. 107(2): p. 359–64.

2. Stupp, R., et al., Effects of radiotherapy with concomitant and adjuvant temozolomide versus radiotherapy alone on survival in glioblastoma in a randomised phase III study: 5-year analysis of the EORTC-NCIC trial. Lancet Oncol, 2009. 10(5): p. 459–66.

3. Phillips, H.S., et al., Molecular subclasses of high-grade glioma predict prognosis, delineate a pattern of disease progression, and resemble stages in neurogenesis. Cancer Cell, 2006. 9(3): p. 157–73.

4. Freije, W.A., et al., Gene expression profiling of gliomas strongly predicts survival. Cancer Res, 2004. 64(18): p. 6503–10.

5. Verhaak, R.G., et al., Integrated genomic analysis identifies clinically relevant subtypes of glioblastoma characterized by abnormalities in PDGFRA, IDH1, EGFR, and NF1. Cancer Cell, 2010. 17(1): p. 98–110.

6. Brennan, C.W., et al., The somatic genomic landscape of glioblastoma. Cell, 2013. 155(2): p. 462–77.

7. Yan, H., et al., IDH1 and IDH2 mutations in gliomas. N Engl J Med, 2009. 360(8): p. 765–73.

8. Noushmehr, H., et al., Identification of a CpG island methylator phenotype that defines a distinct subgroup of glioma. Cancer Cell, 2010. 17(5): p. 510–22.

9. Pearson, J.R.D. and T. Regad, Targeting cellular pathways in glioblastoma multiforme. Signal Transduct Target Ther, 2017. 2: p. 17040.

10. Akhavan, D., et al., De-repression of PDGFRβ transcription promotes acquired resistance to EGFR tyrosine kinase inhibitors in glioblastoma patients. Cancer Discov, 2013. 3(5): p. 534–47.

11. Neftel, C., et al., An Integrative Model of Cellular States, Plasticity, and Genetics for Glioblastoma. Cell, 2019. 178(4): p. 835–849.e21.

12. Bhaduri, A., et al., Outer Radial Glia-like Cancer Stem Cells Contribute to Heterogeneity of Glioblastoma. Cell Stem Cell, 2020. 26(1): p. 48–63.e6.

13. Mootha, V.K., et al., PGC-1alpha-responsive genes involved in oxidative phosphorylation are coordinately downregulated in human diabetes. Nat Genet, 2003. 34(3): p. 267–73.

14. Subramanian, A., et al., Gene set enrichment analysis: a knowledge-based approach for interpreting genome-wide expression profiles. Proc Natl Acad Sci U S A, 2005. 102(43): p. 15545–50.

15. Johnson, S.M., et al., RAS is regulated by the let-7 microRNA family. Cell, 2005. 120(5): p. 635–47.

16. Chiu, S.C., et al., Therapeutic potential of microRNA let-7: tumor suppression or impeding normal stemness. Cell Transplant, 2014. 23(4-5): p. 459–69.

17. Patel, A.P., et al., Single-cell RNA-seq highlights intratumoral heterogeneity in primary glioblastoma. Science, 2014. 344(6190): p. 1396–401.

18. Rollbrocker, B., et al., Amplification of the cyclin-dependent kinase 4 (CDK4) gene is associated with high cdk4 protein levels in glioblastoma multiforme. Acta Neuropathol, 1996. 92(1): p. 70–4.

19. Hua, C., et al., Minichromosome Maintenance (MCM) Family as potential diagnostic and prognostic tumor markers for human gliomas. BMC Cancer, 2014. 14: p. 526.

20. Shen, H., et al., Hypoxia, metabolism, and the circadian clock: new links to overcome radiation resistance in high-grade gliomas. J Exp Clin Cancer Res, 2020. 39(1): p. 129.

21. Giunti, L., et al., A microRNA profile of pediatric glioblastoma: The role of NUCKS1 upregulation. Mol Clin Oncol, 2019. 10(3): p. 331–338.

22. Knudsen, A.M., et al., Expression and prognostic value of the transcription factors EGR1 and EGR3 in gliomas. Sci Rep, 2020. 10(1): p. 9285.

23. Sakakini, N., et al., A Positive Feed-forward Loop Associating EGR1 and PDGFA Promotes Proliferation and Self-renewal in Glioblastoma Stem Cells. J Biol Chem, 2016. 291(20): p. 10684–99.

24. Hemmati, H.D., et al., Cancerous stem cells can arise from pediatric brain tumors. Proc Natl Acad Sci U S A, 2003. 100(25): p. 15178–83.

25. Singh, S.K., et al., Identification of a cancer stem cell in human brain tumors. Cancer Res, 2003. 63(18): p. 5821–8.

26. Galli, R., et al., Isolation and characterization of tumorigenic, stem-like neural precursors from human glioblastoma. Cancer Res, 2004. 64(19): p. 7011–21.

27. Ignatova, T.N., et al., Human cortical glial tumors contain neural stem-like cells expressing astroglial and neuronal markers in vitro. Glia, 2002. 39(3): p. 193–206.

28. Chen, J., et al., A restricted cell population propagates glioblastoma growth after chemotherapy. Nature, 2012. 488(7412): p. 522–6.

29. Bao, S., et al., Glioma stem cells promote radioresistance by preferential activation of the DNA damage response. Nature, 2006. 444(7120): p. 756–60.

30. Guilhamon, P., et al., Meta-analysis of IDH-mutant cancers identifies EBF1 as an interaction partner for TET2. Nat Commun, 2013. 4: p. 2166.

31. Velmeshev, D., et al., Single-cell genomics identifies cell type-specific molecular changes in autism. Science, 2019. 364(6441): p. 685–689.

32. Nowakowski, T.J., et al., Spatiotemporal gene expression trajectories reveal developmental hierarchies of the human cortex. Science, 2017. 358(6368): p. 1318–1323.

33. Gutierrez, L., et al., Post-translational processing of p21ras is two-step and involves carboxyl-methylation and carboxy-terminal proteolysis. EMBO J, 1989. 8(4): p. 1093–8.

34. Ahearn, I.M., et al., NRAS is unique among RAS proteins in requiring ICMT for trafficking to the plasma membrane. Life Sci Alliance, 2021. 4(5).

35. Li, S., et al., RAS/ERK signaling controls proneural genetic programs in cortical development and gliomagenesis. J Neurosci, 2014. 34(6): p. 2169–90.

36. Ricci-Vitiani, L., et al., Tumour vascularization via endothelial differentiation of glioblastoma stem-like cells. Nature, 2010. 468(7325): p. 824–8.

37. Soda, Y., et al., Transdifferentiation of glioblastoma cells into vascular endothelial cells. Proc Natl Acad Sci U S A, 2011. 108(11): p. 4274–80.

38. Cheng, L., et al., Glioblastoma stem cells generate vascular pericytes to support vessel function and tumor growth. Cell, 2013. 153(1): p. 139–52.

39. Horvath, S., et al., Analysis of oncogenic signaling networks in glioblastoma identifies ASPM as a molecular target. Proc Natl Acad Sci U S A, 2006. 103(46): p. 17402–7.

40. Auvergne, R.M., et al., Transcriptional differences between normal and glioma-derived glial progenitor cells identify a core set of dysregulated genes. Cell Rep, 2013. 3(6): p. 2127–41.

41. Park, C., I. Lee, and W.K. Kang, E2F-1 is a critical modulator of cellular senescence in human cancer. Int J Mol Med, 2006. 17(5): p. 715–20.

42. Laine, A., et al., Senescence sensitivity of breast cancer cells is defined by positive feedback loop between CIP2A and E2F1. Cancer Discov, 2013. 3(2): p. 182–97.

43. Wang, L., et al., The Phenotypes of Proliferating Glioblastoma Cells Reside on a Single Axis of Variation. Cancer Discov, 2019. 9(12): p. 1708–1719.

44. Garofano, L., et al., Pathway-based classification of glioblastoma uncovers a mitochondrial subtype with therapeutic vulnerabilities. Nat Cancer, 2021. 2(2): p. 141–156.

45. Laks, D.R., et al., Large-scale assessment of the gliomasphere model system. Neuro Oncol, 2016. 18(10): p. 1367–78.

46. Tirosh, I., et al., Single-cell RNA-seq supports a developmental hierarchy in human oligodendroglioma. Nature, 2016. 539(7628): p. 309–313.

47. Laks, D.R., et al., Neurosphere formation is an independent predictor of clinical outcome in malignant glioma. Stem Cells, 2009. 27(4): p. 980–7.

48. Team, R.C. R: A language and environment for statistical computing. 2020; Available from: https://www.R-project.org/.

