## Supplementary Figures for "Pathway-based approach reveals differential sensitivity of glioblastoma to E2F1 inhibition"

**Supplementary Figure 1. Consensus clustering identifies three clusters for the TCGA dataset.** (A and D) Non-negative matrix factorization (NMF) was performed with ranks ranging from 2 to 6. Various metrics are shown for canonical (A) and oncogenic (D) gene set collections. (B and E) Consensus clustering based on NMF with ranks 2 through 4, showing general structure for each rank in both canonical (B) and oncogenic (E) collections. (C and F) Random Forest classifier was used to validate the robustness of the number of clusters selected. For each analysis, canonical (C) and oncogenic (F), training sets were generated and shown are the results for the correct clustering in the validations sets.

**Supplementary Figure 2. Pathway-based clustering does not overlap with molecular classification.** Principal component analysis plots generated in Figure 2C and 2D were colored by molecular subtype in canonical (left) and oncogenic (right), respectively. Samples for which no subtype was determined are labeled as NA (black). Circle lines represent the normal distribution of the samples in each cluster.

**Supplementary Figure 3. Gliomasphere dataset analysis generates two clusters and TCGA gene ontology analysis of clusters shows differentially enriched terms.** Samples from a 70-sample patient-derived gliomasphere dataset were analyzed using canonical (A) and oncogenic (B) pathways from the GSEA to generate heatmaps based on the enrichment profile of each sample (column) with respect to each gene set (row). (C) Profiles from (A) at the top and (B) at the bottom were used to generate PCA plots labeled by color and shape of each cluster. Circle lines represent the normal distribution of the samples in each cluster. (D) TCGA samples were reanalyzed utilizing the gene lists generated in Figure 4 to obtain enrichment profiles for each sample (not shown) and the corresponding shown PCA plot. The arrows show the contribution of each gene list to a particular direction in the plot. (E) The three clusters generated for the TCGA dataset were analyzed using differential expression analysis and then analyzed for the enrichment of gene ontology terms in each cluster. The plot shows the terms most highly enriched in a particular cluster compared with the other two. The expression of marker genes in each cluster is shown for every sample.

**Supplementary Figure 4. Correlation between gene lists in TCGA and gliomasphere datasets.** The generated gene lists were used to obtain scores for each sample in both TCGA (top) and GS (bottom) datasets. Correlations between these scores in all samples for either dataset are shown, where the presence of a circle represents significant correlation and the size and color depict the strength of the correlation. Boxes mark groups of gene lists strongly correlated (based on hierarchical clustering).

**Supplementary Figure 5. Pathway-based clustering in single cell dataset reveals low intratumoral heterogeneity.** Samples from a single cell RNA-seq database were analyzed using either canonical pathways (left) or oncogenic pathways (right) from the Gene Set Enrichment Analysis to generate heatmaps based on the enrichment profile of each sample (column) with respect to each gene set (row) in both collections. Colors at the top represent the sample to which each cell belongs (MGH26 = grey, MGH264 = magenta, MGH28 = cyan, MGH29 = orange, MGH30 = green, MGH31 = purple; grey and magenta overlap the most because MGH26 and MGH264 are different analyses from the same sample).

**Supplementary Figure 6. Gene lists differentially correlate with cell cycle and stemness signatures.** (A and B) Gene lists were used to obtain scores for each cell in the scRNA-seq dataset [17]. Similarly, stemness and cell cycle score scores were generated using the signatures reported in the same paper. For each cell, we plotted the scores for cell cycle (A) or stemness (B) signatures against the scores for all the gene lists. Cells are colored by tumor of origin and p-values and correlation coefficients are shown for each plot.

**Supplementary Figure 7. Gene lists differentially correlate with cellular states and cell specific markers.** Scores generated for each cell in Supplementary Figure 5 using gene lists were correlated with scores for cellular states ([11]) and specific cell type markers in development and adult brain ([31, 32]). The presence of a circle represents significant correlation, and the size and color depict the intensity of the correlation. Boxes mark groups of gene lists strongly correlated (based on hierarchical clustering).

**Supplementary Figure 8. E2F1-enriched samples have high enrichment scores for DNA repair gene sets.** Samples in the gliomasphere dataset from both clusters (determined in Figure 4) were evaluated for their enrichment of DNA repair gene sets from the canonical pathway collection.

**Supplementary Table 1** – Gene lists for each PC and direction

**Supplementary Table 2** – Gene sets enriched in cell cycle cluster for scRNA-seq

**Supplementary Table 3** – Summary from enrichR and IPA for non-cell cycle clusters

**Supplementary Table 4** – WCGNA GO enriched terms for modules associated with both clusters

### Supplementary Figure 1

A

#### Canonical Pathways

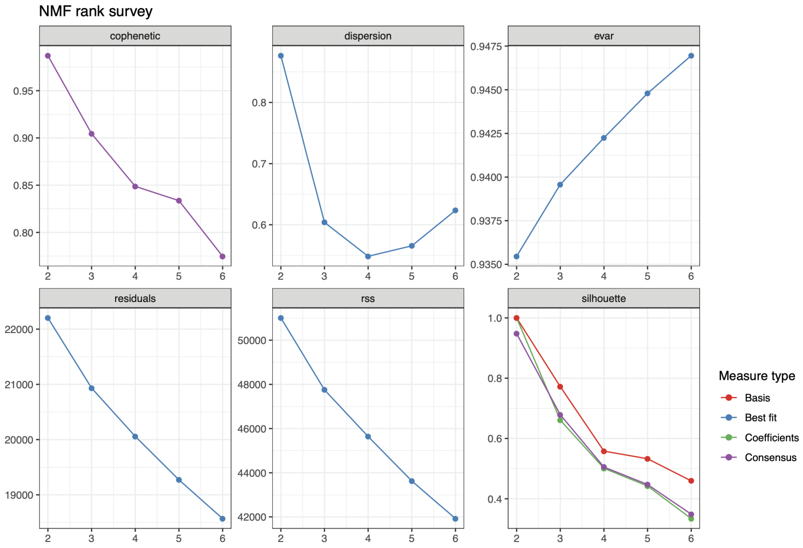


B

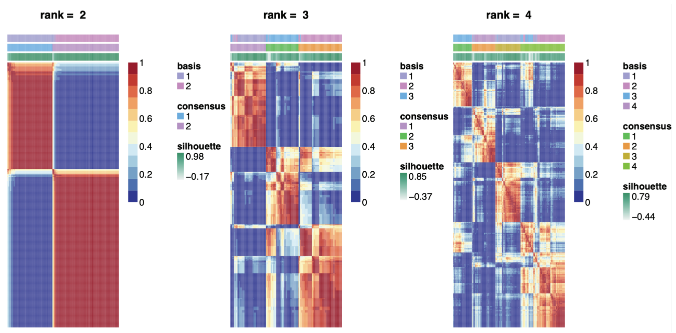


C

```
> predValidC2 <- predict(modelC2, ValidSetC2, type = "class")
> mean(predValidC2 == ValidSetC2$cluster)
[1] 0.9320988
> table(predValidC2,ValidSetC2$cluster)

predValidC2    1     2     3
          1   28     2     0
          2    3    58     4
          3    2     0    65
```

D

#### Oncogenic Pathways

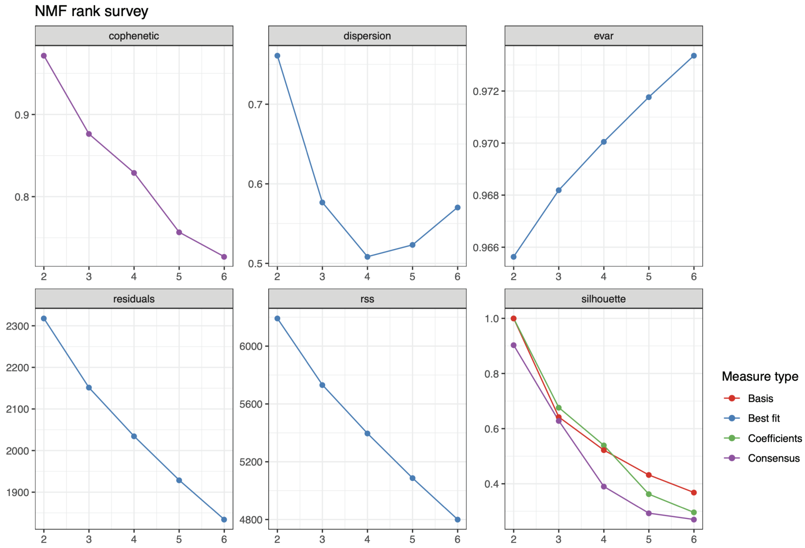


E

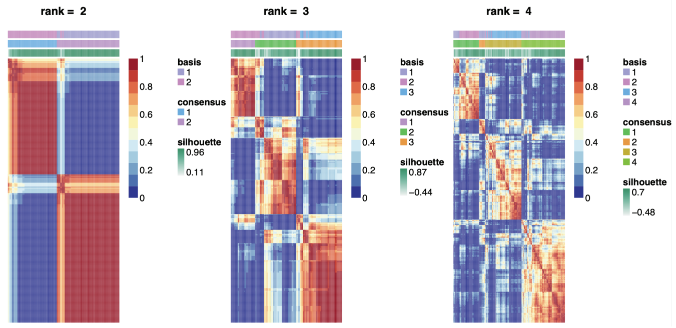


F

```
> predValidC6 <- predict(modelC6, ValidSetC6, type = "class")
> mean(predValidC6 == ValidSetC6$Cluster)
[1] 0.9753086
> table(predValidC6,ValidSetC6$Cluster)

predValidC6    1     2     3
          1   34     1     0
          2    1    68     2
          3    2     0    56
```

### Supplementary Figure 2

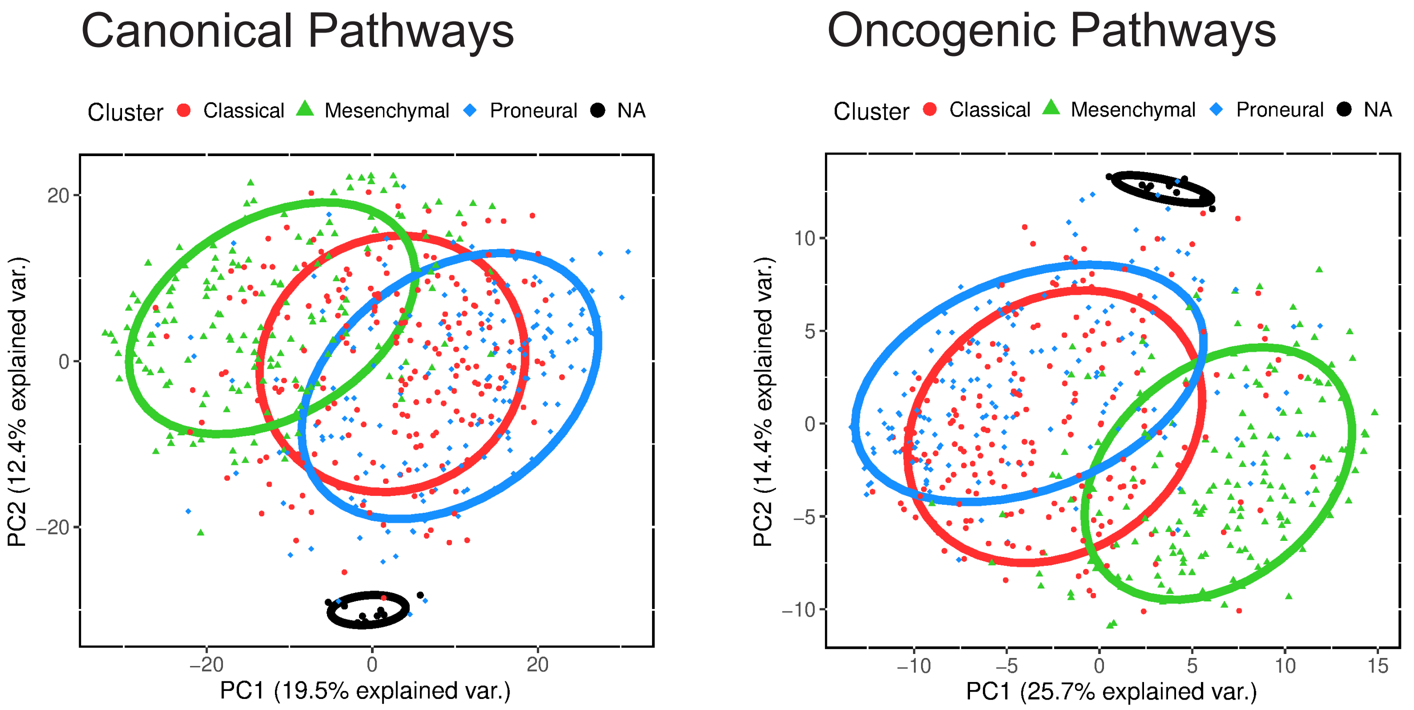

### Supplementary Figure 3

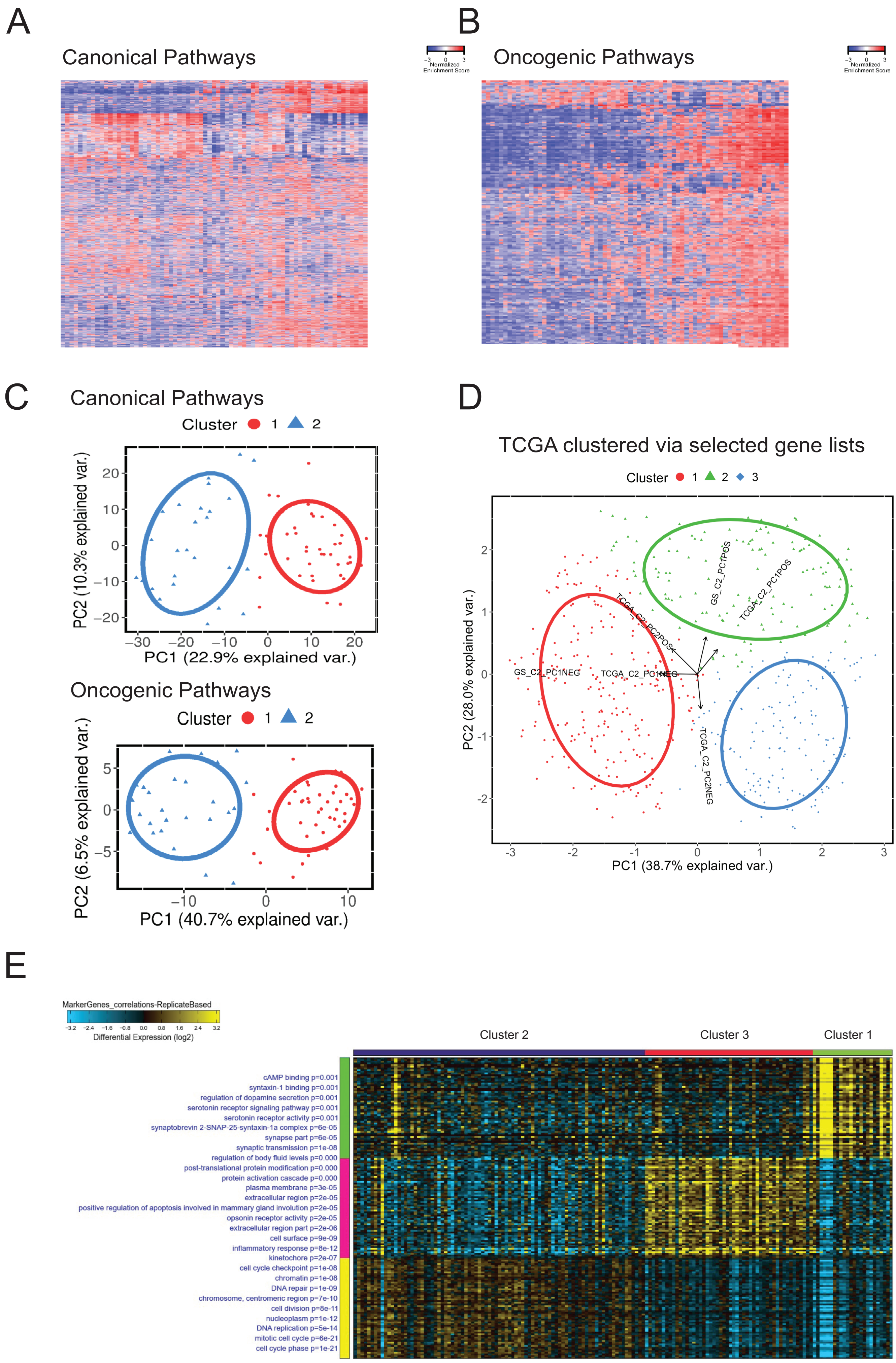

### Supplementary Figure 4

#### TCGA dataset

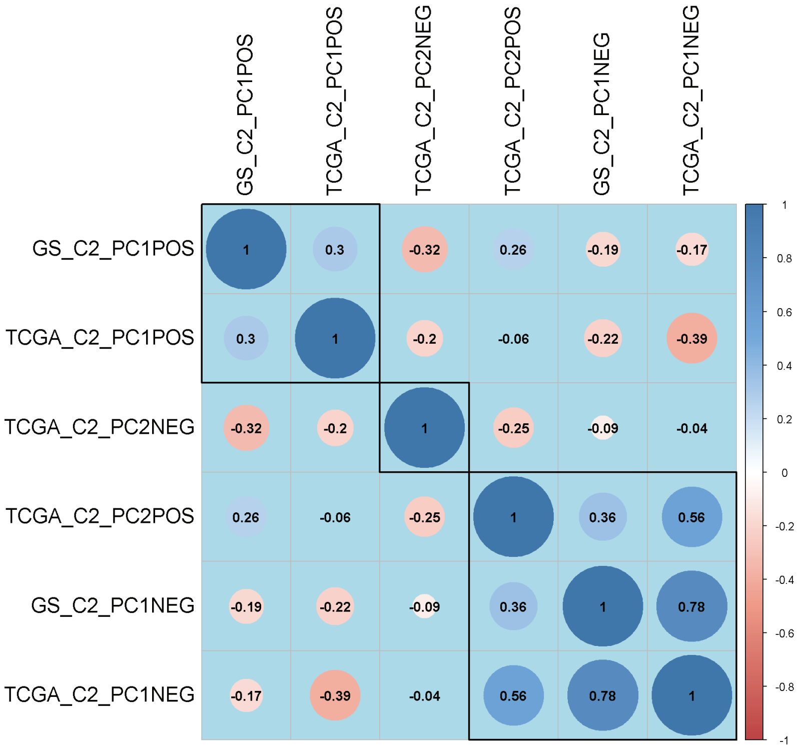


#### Gliomasphere dataset

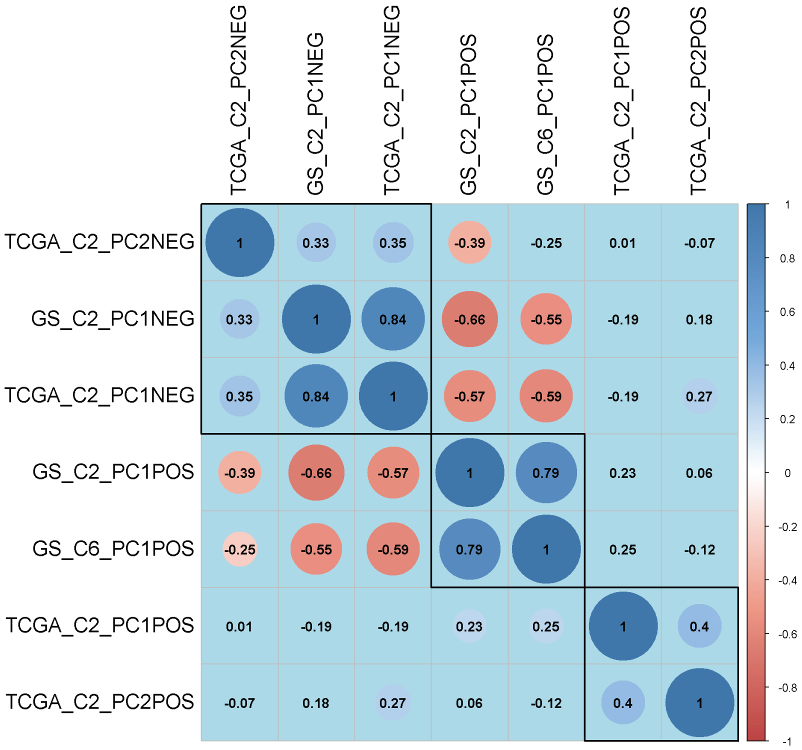

### Supplementary Figure 5

#### Canonical Pathways

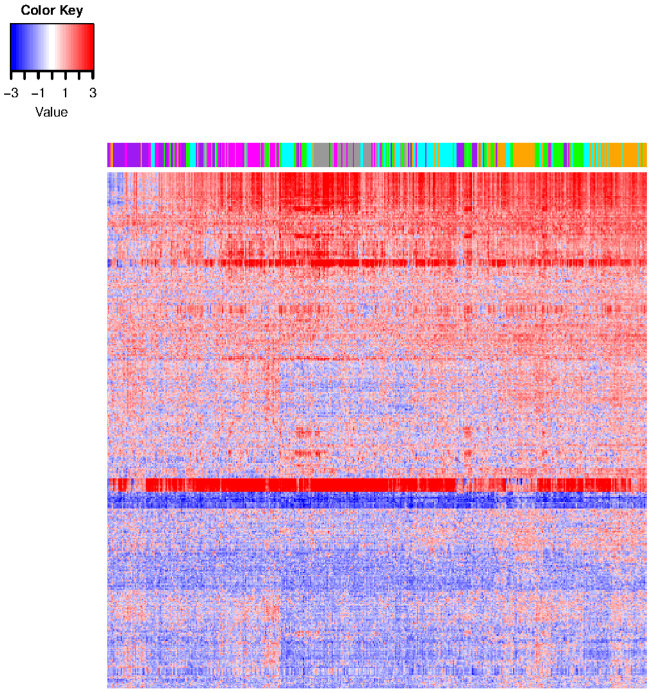


#### Oncogenic Pathways

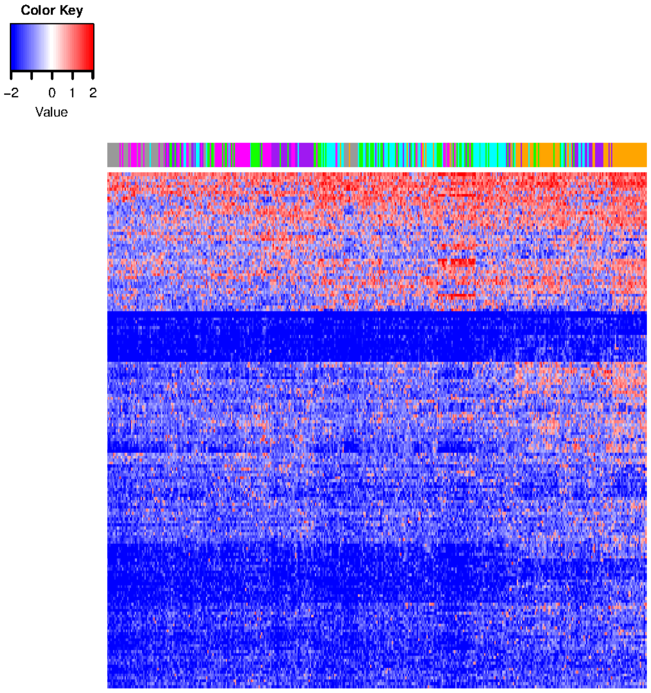

### Supplementary Figure 6

A

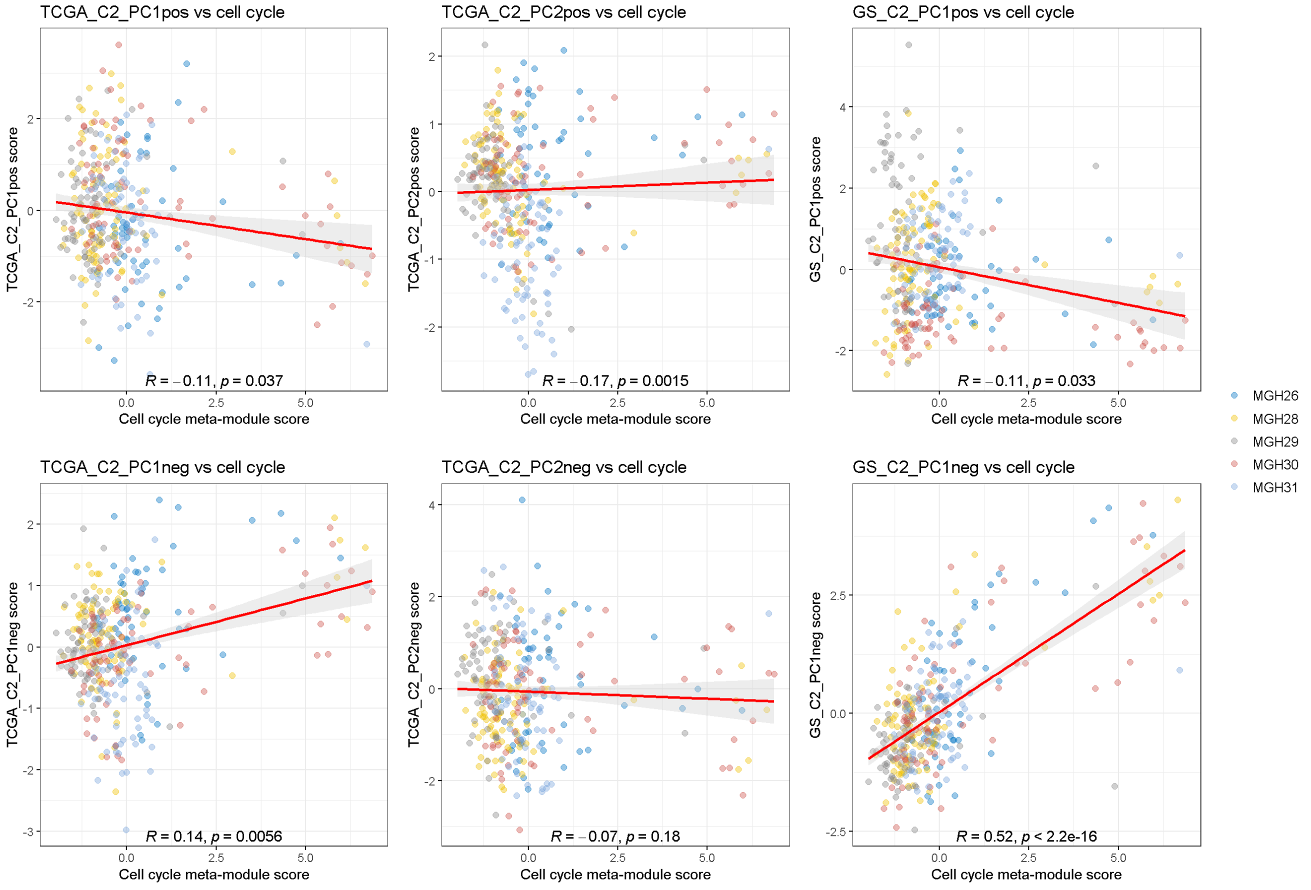


B

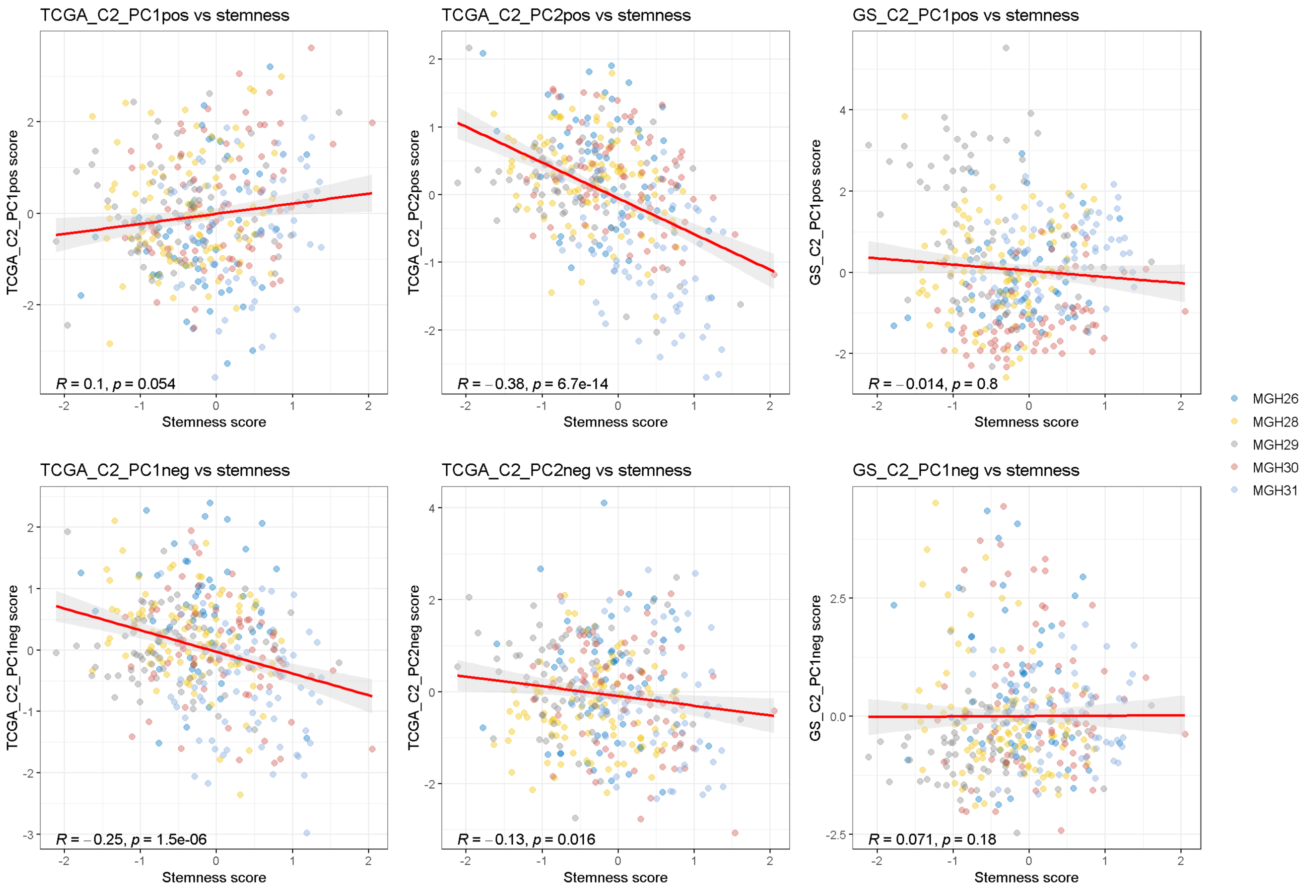

### Supplementary Figure 7

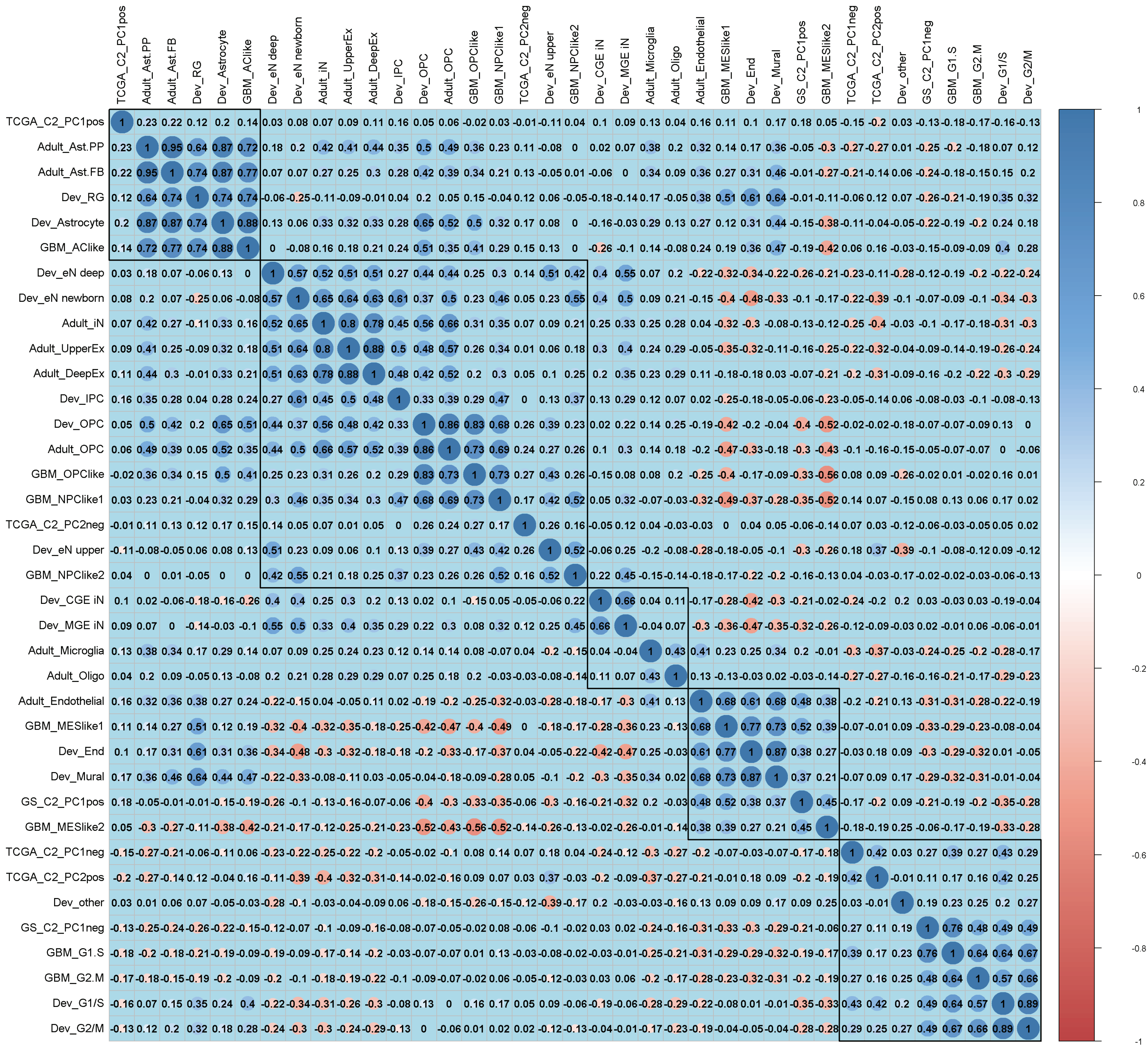

### Supplementary Figure 8

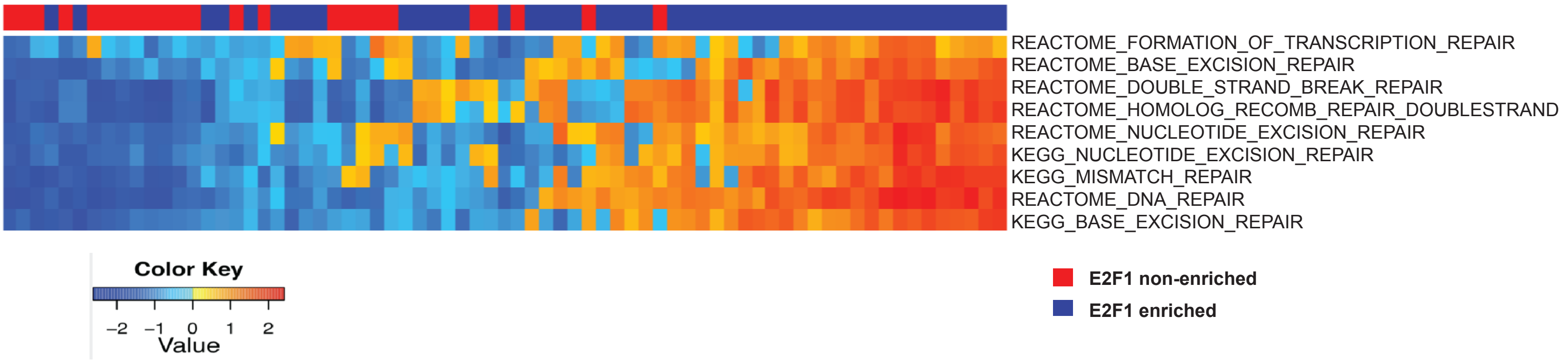
